# Hierarchical TBX6-FOXC Regulatory Logic Governs Human Trunk Mesoderm Diversification

**DOI:** 10.64898/2026.03.17.712295

**Authors:** John-Poul Ng-Blichfeldt, Rosalind Drummond, David Lando, Samina Kausar, Melania Barile, Anna Philpott

## Abstract

Mesoderm forms during gastrulation and diversifies to generate many embryonic tissues, yet the regulatory logic governing early human mesodermal lineage decisions remains poorly defined. We use human iPSC-derived 3D trunk-like structures (hTLS) to model lineage specification during early post-gastrulation development, a stage experimentally inaccessible in humans. Developing hTLS generate neural and mesodermal lineages and recapitulate progressive mesodermal diversification into paraxial (somitic) and intermediate (renal) identities. Using genetic perturbations and temporally controlled transcription factor activation, we identify duration-dependent TBX6 activity as a critical determinant of mesodermal diversification. Downstream, the forkhead box transcription factors FOXC1 and FOXC2 stabilise somitic identity and are required for sclerotome differentiation. These findings reveal a duration-dependent hierarchical regulatory logic in early human mesoderm, in which competence for diversification is established before downstream lineage identities are stabilised, positioning hTLS as a powerful platform to experimentally dissect human developmental programs.

**Highlights:**

- Human trunk-like structures model mesodermal diversification during early post-gastrulation human trunk development
- TBX6 duration acts as a temporal gate controlling mesodermal fate potential
- FOXC1/2 function downstream of TBX6 to lock in somitic identity and restrict alternative fates
- DegTrace enables lineage tracing following transient transcription factor activation

## Introduction

How transient developmental states are converted into stable lineage identities remains a fundamental problem in developmental biology. Embryonic cell identity begins to diversify prior to and during gastrulation as epiblast cells progressively generate the three embryonic germ layers, with cells that ingress through the primitive streak forming mesoderm and endoderm while non-ingressing cells adopt ectodermal identity ^1,2^. Mesoderm further diversifies into distinct lineages that form the primordia of a wide range of tissues, with specific mesodermal identities arising from discrete positions along the rostral-caudal axis of the streak ^3^. Emerging from the prospective trunk region of the embryo, paraxial mesoderm undergoes segmentation into somites, transient epithelial structures that further subdivide into sclerotome that forms vertebrae, and dermomyotome that produces skeletal muscle ^4^. Adjacent to paraxial mesoderm, intermediate mesoderm gives rise to the urogenital system, including the tubular epithelial components of the kidneys and gonads ^5,6^. Despite its fundamental importance, the regulatory logic by which early mesoderm diversifies into distinct lineages, including paraxial and intermediate mesoderm and their derivatives, remains incompletely understood.

Classic fate-mapping and grafting experiments in avian and mouse embryos demonstrated that cells of the primitive streak represent a transient multipotent state characterised by broad equivalence in lineage competence, with positional cues within the streak directing specific mesodermal identities ^7–13^. These positional cues are established by signalling gradients that both pattern the embryonic axes and bias lineage specification: Wnt activity posteriorises streak identity and promotes mesodermal differentiation, whereas retinoic acid (RA) antagonises Wnt/FGF signalling to permit differentiation of trunk progenitors while also contributing to posterior neural patterning ^14,15^. However, downstream of local signalling environments, how cell-intrinsic transcription factor dynamics convert transient multipotent states into stable lineage-specific mesodermal identities remains poorly defined.

Recent comparative studies have highlighted divergence in developmental mechanisms between mouse and human mesodermal derivatives, including differences in morphogenesis and patterning of the kidney ^16–19^ and in the tempo and signalling mechanisms underlying somitogenesis ^20,21^. However, direct investigation of these early developmental events in humans is constrained by technical and ethical limitations ^22^. Advances in single-cell transcriptomics have provided valuable insights into transcriptional states associated with early lineage diversification in human embryos, yet these stages remain experimentally inaccessible *in vivo* ^23–25^. Recently, the development of human pluripotent stem cell–based 3D differentiation systems has begun to enable functional investigations of early human developmental programs by modelling key aspects of embryonic development *in vitro* ^26–34^.

Here, we employ human induced pluripotent stem cell (iPSC)-derived trunk-like structures (hTLS) to investigate the molecular mechanisms governing early diversification of paraxial (somitic) and intermediate (renal) mesoderm lineages during early post-gastrulation human trunk development. By combining temporal control of transcription factor activity with genetic perturbations, we uncover a previously unknown regulatory program governing mesodermal diversification. We find that the duration of upstream TBX6 activity establishes a transient window of multipotent competence, which is subsequently stabilised by the downstream transcription factors FOXC1 and FOXC2 to drive lineage commitment. Together, these findings reveal a duration-dependent hierarchical regulatory logic governing human mesoderm diversification and demonstrate the utility of hTLS as a tractable experimental system to dissect early human developmental programs.

## Results

### Human trunk-like structures (hTLS) model progressive mesodermal diversification

To investigate how trunk mesodermal lineages emerge and diversify during early post-gastrulation human development, we adapted published protocols to generate 3D trunk-like structures (hTLS) from human iPSCs. In this system, iPSCs are exposed to a transient pulse of high Wnt signalling, followed by reduced Wnt signalling combined with RA, prior to aggregation ^27^. Under these signalling conditions, neural and mesodermal identities emerge concomitantly, recapitulating lineage diversification characteristic of posterior (post-cranial) embryonic development (Figure 1A) ^27^. Differentiating aggregates embedded in Matrigel elongated and generated SOX2^+^ neural and SOX2^-^non-neural tissues in parallel (Figures 1B and S1A,A’).

**Figure 1.**
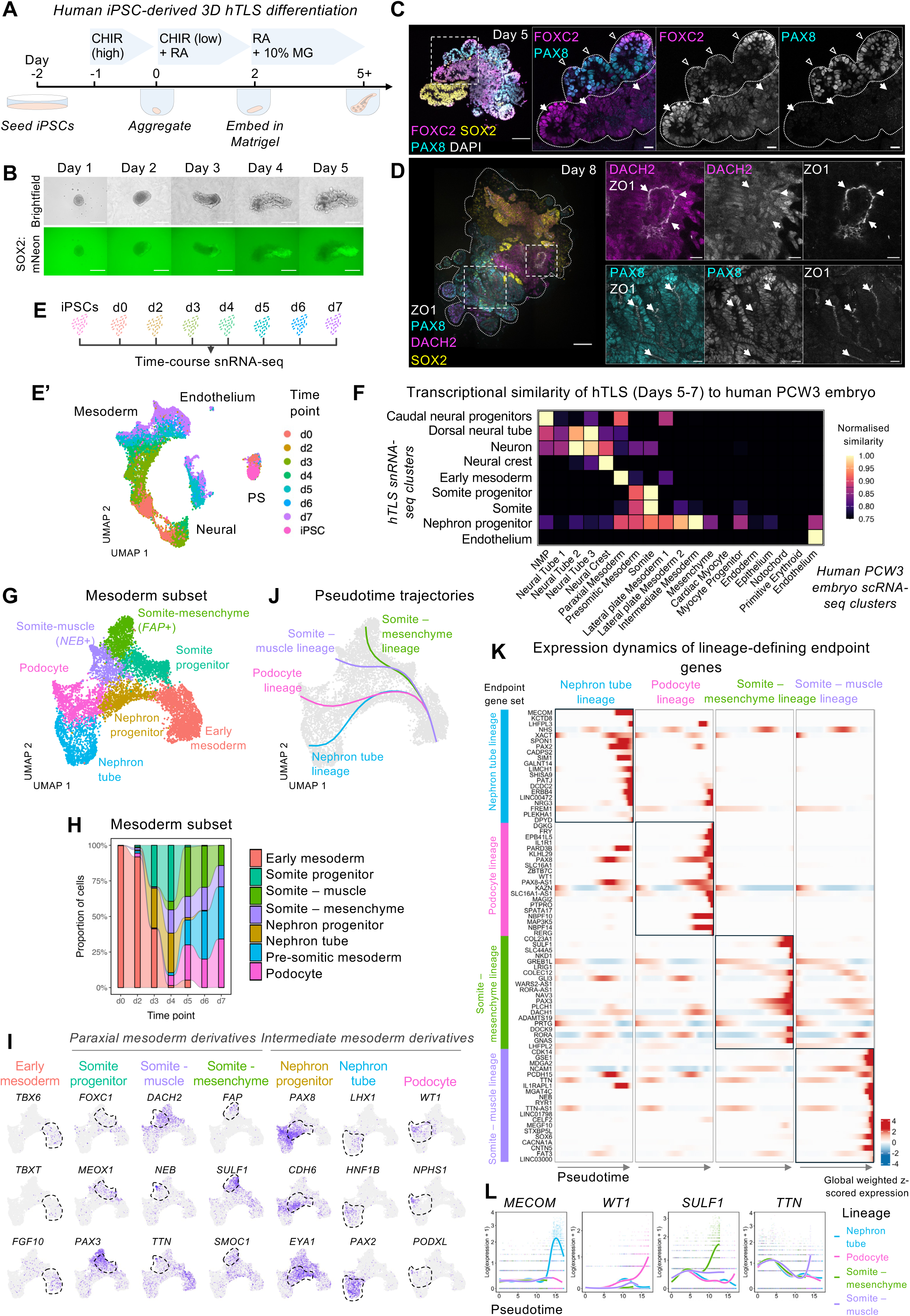
Human trunk-like structures recapitulate progressive mesodermal diversification. (A) Schematic of the protocol used to generate human trunk-like structures (hTLS) from induced pluripotent stem cells (iPSCs), adapted from ^27^. (B) Brightfield and corresponding green fluorescence images of hTLS generated from SOX2:mNeon iPSCs from day 1 to day 5 of the protocol, showing expansion of mNeon^+^ and mNeon^-^ domains. Scale bars = 300 µm. (C and D) Immunofluorescence of hTLS showing (C) FOXC2 (magenta), SOX2 (yellow), PAX8 (cyan) expression at day 5, with DAPI as nuclear counterstain (grey), or (D) DACH2 (magenta), SOX2 (yellow), PAX8 (cyan), and ZO1 (grey) expression at day 8. hTLS are outlined with dotted white lines. Scale bars: main, 100 µm; insets, 20 µm. (E) Schematic of the hTLS single-nucleus RNA-sequencing (snRNA-seq) strategy. (E’) UMAP representation of 13,618 hTLS cells coloured by time point, with major lineages annotated (see Supplemental Figure 1G,H). PS, pluripotent stem cells. (F) Heatmap showing transcriptional similarity between clusters identified from hTLS snRNA-seq (days 5-7) and clusters from post-conception week 3 (PCW3) human embryo scRNA-seq data ^24^. (G) UMAP representation of 8,088 cells of hTLS mesodermal cells identified from the snRNA-seq dataset (see Supplemental Figure S1M, N), coloured by unsupervised clustering and annotated based on marker expression. (H) Alluvial plot showing the proportion of hTLS mesodermal cells at each time point assigned to clusters defined in (G). (I) Expression of representative marker genes for clusters in (G) overlaid on UMAP plots. (J) UMAP representation of hTLS mesodermal cells (as in G) with pseudotime trajectories inferred using Slingshot ^36^. (K) Within-lineage standardized expression dynamics of the top 20 genes defining each terminal state along the pseudotime trajectories shown in (J), identified using tradeSeq endpoint differential expression ^37^. Genes are ranked by the strength and consistency of their differential expression at lineage endpoints. Heatmaps show z-scored fitted expression along pseudotime within each lineage (red, higher; blue, lower). (L) Expression of representative genes identified in (K) (*MECOM, WT1, SULF1* and *TTN*) plotted along pseudotime for each lineage. Individual cells are shown as dots, with fitted expression curves shown as lines and coloured by lineage. See also Figure S1.

At day 5, SOX2^+^ structures exhibited lumens and expressed PAX6, consistent with neural tube-like identity (Figure S1B) ^35^. Surrounding SOX2^-^ tissue expressed the paraxial mesoderm marker forkhead box C2 (FOXC2), and the nephron progenitor marker PAX8, indicating the concurrent emergence of both paraxial (somitic) and intermediate (renal) mesodermal identities (Figures 1C and S1B). By day 8, hTLS contained somitic (DACH2^+^) and renal (PAX8^+^) epithelial structures with apical-basal polarity, indicated by the apical localisation of ZO1 (Figure 1D). Renal epithelial structures expressed CDH1 (E-Cadherin/ECAD), hepatocyte nuclear factor beta (HNF1B) and PAX8, consistent with nephron tubular identity (Figure S1C). Similar neural, somitic, and renal lineages were observed in hTLS generated from an independent iPSC line (IMR-90-4), demonstrating protocol robustness across different genetic backgrounds (Figure S1D).

To define the transcriptional programs underlying lineage emergence in hTLS, we performed single nucleus RNA-sequencing (snRNA-seq) across a time course from day 0 to day 7, including undifferentiated iPSCs as a control (Figures 1E,E’ and S1E-I). Unsupervised clustering revealed *MEOX1*^+^ somitic mesoderm, *PAX2*^+^ intermediate mesoderm-derived kidney progenitors and their derivatives (*HNF1B*^+^ nephron tube and *NPHS1*^+^ podocytes), as well as neural tube-derived *MAP2*^+^ neurons and *SOX10*^+^ neural crest-like populations (Figures S1G-I). By immunofluorescence, cells adjacent to neural tube-like structures exhibited migratory morphology and were SOX9^+^, consistent with a neural crest-like identity (Figure S1J). *PECAM-1*^+^ endothelial cells were also identified by both snRNA-seq and immunofluorescence (Figures S1G-I and S1K).

To assess the developmental correspondence of hTLS to human embryogenesis *in vivo*, we compared the hTLS snRNA-seq dataset to reference atlases spanning zygote to post-conception week 2 (PCW2/CS7) ^25^, and to an independent PCW3/CS10 human embryo dataset ^24^. hTLS cells from days 0-4 mapped predominantly to CS7-stage embryonic populations (Figures S1G and SL-L’’), whereas cells from days 5-7 showed strong correspondence to PCW3 neural tube, somite, intermediate mesoderm and endothelial lineages, while lacking notochord, endoderm, cardiac and erythroid populations (Figures 1F and S1G,H). These analyses support the use of hTLS as an experimentally tractable model of early post-gastrulation human trunk development, capturing neural tube, and paraxial (somitic) and intermediate (renal) mesodermal diversification.

To examine mesodermal diversification in greater detail, we isolated mesodermal cells in the hTLS snRNA-seq dataset and performed secondary unsupervised clustering (Figures 1G and S1M-P). Early time points were dominated by a single mesodermal population enriched for *TBX6*, *FGF10* and *PDGFRA* (Early mesoderm; Figures 1H and S1P). From day 3 onward, additional clusters emerged corresponding to nephron progenitors (*PAX8*^+^ *CDH6*^+^) and somite progenitors (*MEOX1*^+^ *PAX3*^+^), followed by diversification into nephron derivatives, alongside dorsal somite (*PAX3*^+^ *DACH2*^+^)-derived muscle (*NEB*^+^) and mesenchymal (*FAP*^+^) populations at later time points (Figures 1G,H and S1P). These data indicate that hTLS recapitulate progressive mesodermal diversification into paraxial (somitic) and intermediate (renal) mesoderm derivatives, and therefore enable detailed investigation of mechanisms controlling their divergence.

To reconstruct developmental trajectories underlying mesodermal diversification, we performed pseudotime inference using Slingshot ^36^, identifying four major trajectories corresponding to nephron tube, podocytes, somite-muscle and somite-mesenchymal lineages (Figure 1J). While cluster identities were used to define the overall trajectory structure, gene expression dynamics were analysed along pseudotime at single-cell resolution. Using tradeSeq ^37^, we identified lineage-specific transcriptional programs associated with terminal states along each trajectory (Figures 1K,L). Nephron tube and podocyte lineages exhibited distinct renal epithelial (*PAX2*, *PATJ*) and glomerular (*WT1*, *MAGI2*) gene signatures, whereas somite-muscle and somite-mesenchymal lineages were marked by sarcomeric (*TTN*, *NEB*) or extracellular matrix-associated (*COL23A1*, *SULF1*) gene expression. Together, these analyses establish hTLS as a system in which trunk mesodermal lineages progressively diversify along transcriptionally distinct trajectories, providing a foundation for mechanistic interrogation of human trunk mesoderm specification and diversification.

### TBX6 is required for establishment of a transient multipotent mesodermal competence state in hTLS

To identify factors regulating the initial establishment of mesodermal multipotency in hTLS, we compared the earliest emerging mesodermal and neural populations within the hTLS snRNA-seq dataset. Differential expression analysis identified 27 transcription factors enriched in early mesoderm, with the T-box transcription factor *TBX6* as one of the most highly enriched (Figures 2A,A’). Although TBX6 is classically associated with somitic mesoderm specification, recent work has suggested that its expression precedes overt lineage diversification ^38,39^, raising the possibility that TBX6 functions more broadly to establish mesodermal competence. We therefore examined the temporal dynamics of TBX6 expression and its functional requirement during early hTLS differentiation.

**Figure 2.**
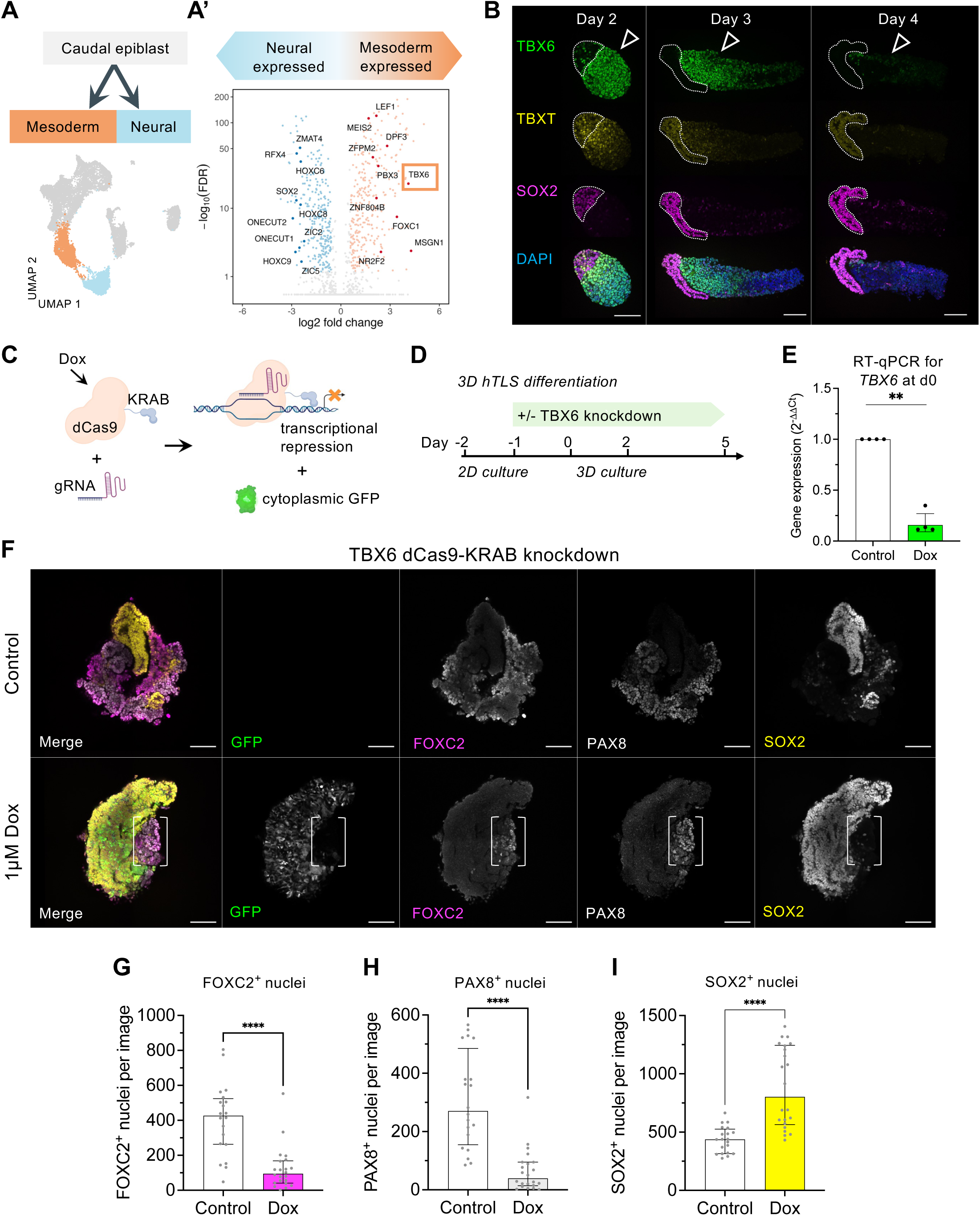
TBX6 defines early mesoderm and is critical for its diversification. (A) Schematic of bifurcation of caudal epiblast cells into mesodermal and neural-fated cells (coloured in UMAP representation of 13,618 hTLS cells underneath). (A’) Genes differentially expressed between mesoderm and neural clusters (as in A). Genes with adjusted p-value < 0.05 and |log2 fold change| > 0.58 (±1.5-fold) are highlighted (red, upregulated; blue, downregulated). Genes encoding the top 10 up- and downregulated transcription factors, ranked by log2 fold change, are shown in darker colours and annotated. (B) Immunofluorescence showing TBX6 (green), TBXT/Brachyury (yellow), and SOX2 (magenta) expression in hTLS at days 2, 3 and 4, with nuclei labelled with DAPI (blue). Putative neuromesodermal progenitors (NMP)-like cells (SOX2^+^, TBXT^+^, TBX6^-^) are outlined with a dotted line. Arrowhead indicates TBX6^+^ mesodermal domain showing progressive loss of TBX6 expression. Scale bars, 100 µm. (C) Schematic of the dCas9-KRAB CRISPR interference (CRISPRi) gene knockdown system ^40^. Created with BioRender.com. (D) Schematic of experimental protocol to suppress *TBX6* expression in hTLS. (E) RT-qPCR for *TBX6* in TBX6-dCas9-KRAB iPSCs differentiated in the hTLS protocol and harvested after 24 hours, either untreated (control) or treated with Dox (1 µM) from day -1. Statistical significance was assessed using two-way ANOVA on log-transformed values, with treatment as the column factor and differentiation batch as the row factor; p value corresponds to the main effect of treatment (** p < 0.01). Data are plotted as 2^-ΔΔCt^ for visualisation; bars represent mean ± SEM; dots represent n = 4 independent differentiation batches per group. (F) Immunofluorescence of GFP (green), SOX2 (yellow), FOXC2 (magenta), or PAX8 (grey) expression in day 5 hTLS generated from TBX6-dCas9-KRAB iPSCs, either untreated (control, top) or treated with Dox from day -1 (1 µM, bottom). White brackets indicate SOX2^-^, FOXC2^+^, PAX8^+^ cells confined to a GFP^-^ region. Scale bars, 100 µm. (G-I) Quantification of number of nuclei per confocal image positive for (G) FOXC2, (H) PAX8, or (I) SOX2, in day 5 TBX6-dCas9-KRAB hTLS either untreated (control, white bars; n=20 hTLS across 2 independent batches) or treated with Dox from day -1 (1 µM, coloured bars; n=22 hTLS across 2 independent batches). Dots represent individual hTLS; bars represent median ± IQR; Mann-Whitney test. See also Figure S2.

We observed by immunofluorescence that TBX6 was expressed at low levels and heterogeneously in iPSCs after 24 hours of differentiation (day 0), whereas the primitive streak marker TBXT/Brachyury was broadly expressed (Figures S2A,B). Notably, TBX6 expression was restricted to a subset of TBXT^+^ cells, indicating that TBX6 marks a subpopulation within the broader TBXT^+^ mesodermal compartment. A subset of TBXT^+^ cells also co-expressed SOX2 at early stages (Figures 2B and S2B), consistent with neuromesodermal progenitor (NMP)-like states that may contribute to both neural and mesodermal lineages in this system. By day 2 in aggregated hTLS, TBX6 and SOX2 marked distinct, non-overlapping cell populations, with TBX6 expression declining at day 3 and largely depleted by day 4 (Figure 2B). Time course RT-qPCR analyses revealed that *TBX6* induction preceded markers of both somitic mesoderm (*FOXC2*, *MEOX1*, and *PAX3)* and intermediate (renal) mesoderm (*OSR1*, *PAX2*, *PAX8*; Figures S2C,D). Thus, consistent with observations in mouse embryos *in vivo* ^38^, TBX6 is transiently expressed in mesoderm prior to lineage diversification in hTLS, suggesting a broader role in establishing mesodermal competence.

We next investigated the role of TBX6 in mesodermal emergence. We engineered iPSCs with a CRISPR interference (CRISPRi) system to enable inducible suppression of TBX6 expression (Figures 2C,D) ^40^ . In this system, a catalytically dead Cas9 (dCas9) fused to the Krüppel-associated box (KRAB) repressor domain is targeted to the *TBX6* locus under the control of Doxycycline (Dox), with cytoplasmic GFP as a readout for dCas9 expression (TBX6-dCas9-KRAB). Addition of Dox at the onset of differentiation (day -1) resulted in ∼82% knockdown of *TBX6* expression at day 0 compared to untreated cells (p = 0.0042; Figure 2E). *TBX6* knockdown was associated with decreased expression of the early mesodermal markers *TBXT* (p = 0.011) and *MSGN1* (p = 0.0031; Figure S2E). Depletion of TBX6 protein after Dox treatment was confirmed by immunofluorescence (Figure S2F). Next, TBX6-dCas9-KRAB iPSCs were differentiated to hTLS and harvested at day 5 of the protocol (Figure 2D). Control hTLS did not express GFP, indicating lack of knockdown cassette expression, and exhibited somitic (FOXC2^+^), renal (PAX8^+^) and neural (SOX2^+^) tissue as observed previously (Figure 2F). By contrast, Dox-treated TBX6-dCas9-KRAB hTLS showed heterogeneous GFP expression with a reduction in both FOXC2^+^ and PAX8^+^ tissue and an expanded SOX2^+^ domain. Remaining FOXC2^+^PAX8^+^SOX2^-^ cells were GFP^-^ and thus did not express the knockdown cassette, indicating that renal and somitic mesodermal differentiation was mutually exclusive with TBX6 depletion (Figure 2F). Quantification revealed a 78% decrease in the number of FOXC2^+^ cells per confocal image (p < 0.0001; Figure 2G), and an 85% decrease in PAX8^+^ cells (p < 0.0001; Figure 2H), with a corresponding 84% increase in SOX2^+^ cells (p < 0.0001; Figure 2I). Similar results were obtained with an independent iPSC line with a distinct gRNA to target TBX6 (Figure S2G). These data indicate that TBX6 is necessary for establishment of a multipotent mesodermal population in hTLS, from which both somitic and renal lineages emerge.

### Acute TBX6 expression promotes mesodermal transcriptional progression without commitment to a specific lineage

TBX6 is transiently and heterogeneously expressed early in hTLS differentiation (Figures 2B and S2B). This raised the question of whether TBX6 is permissive for mesodermal diversification or actively instructs mesodermal progression. To address this, we engineered a piggyBac-insertable, Dox-inducible TBX6 overexpression system with cytoplasmic mCherry as a fluorescent readout of transgene induction, to more uniformly elevate TBX6 expression in the earliest stages of hTLS differentiation (PB-Dox-TBX6; Figures 3A-C).

**Figure 3.**
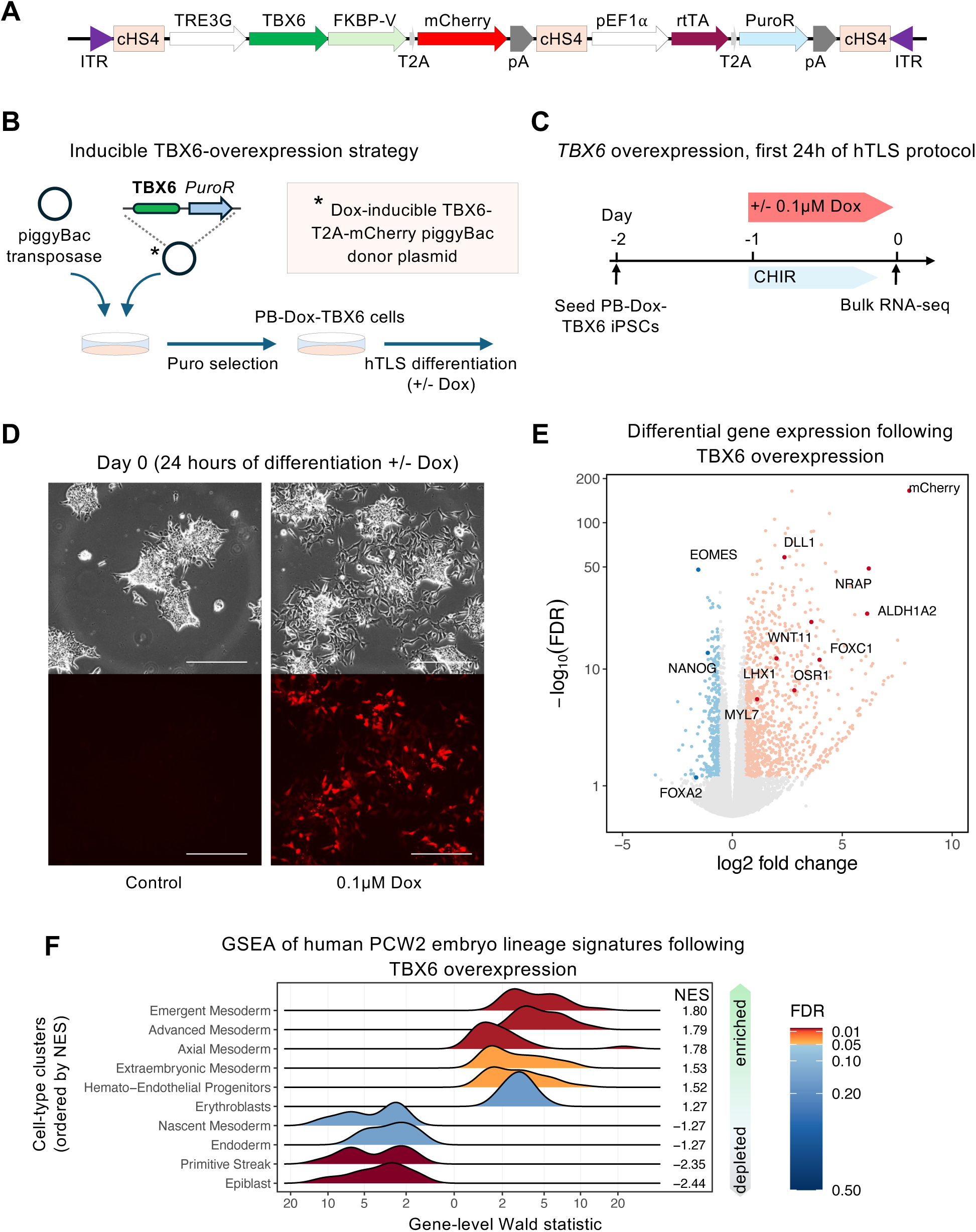
TBX6 drives transcriptional progression toward advanced mesodermal fates. (A) Schematic of a piggyBac-insertable construct for doxycycline (Dox)-inducible overexpression of TBX6-FKBP-T2A-mCherry. (B) Inducible TBX6 transgene overexpression strategy. A piggyBac donor plasmid encoding a Dox-inducible TBX6-FKBP-T2A-mCherry cassette with puromycin resistance was integrated into iPSCs using piggyBac transposase. Following selection, edited cells were differentiated to hTLS with or without Dox to induce transgene expression. (C) Experimental design for TBX6 overexpression during the first 24 hours of the hTLS protocol in piggyBac-edited iPSCs (PB-Dox-TBX6), followed by bulk RNA-seq. (D) Brightfield and corresponding red fluorescence images of PB-Dox-TBX6 cells 24 hours after initiation of the hTLS protocol, either without Dox (control) or with Dox (0.1 µM). Scale bars, 300 µm. (E) Volcano plot of differential gene expression following 24 h of TBX6 overexpression versus control (n = 4 independent differentiation batches per condition). Genes with adjusted p-value < 0.05 and |log2 fold change| > 0.58 (±1.5-fold) are highlighted (red, upregulated; blue, downregulated). Selected genes associated with alternative lineages are annotated and shown in darker colour. (Somitic: *FOXC1, DLL1, ALDH1A2*; intermediate (renal) mesoderm: *OSR1, LHX1*; cardiac/muscle lineages: *MYL7, NRAP*; pluripotency: *NANOG*; mesendoderm/endoderm: *EOMES, FOXA2*). (F) Ridgeplots of gene set enrichment analysis (GSEA) of genes following TBX6 overexpression (Dox versus control), ranked by the DESeq2 Wald statistic ^72^, and tested against the top 50 marker genes for each annotated PCW2 human embryo scRNA-seq population from Tyser et al. ^23^. Ridgeplots show the distribution of DESeq2 Wald statistics for genes within each population marker set along the ranked gene list; populations are ordered by normalised enrichment score (NES), and colour denotes FDR-adjusted p-values. ITR, inverted terminal repeat; TRE3G, third-generation tetracycline-responsive promoter; cHS4, chicken hypersensitive site 4 insulator; pA, polyadenylation signal; PuroR, puromycin resistance gene; GSEA, gene set enrichment analysis; PCW2, post-conception week 2. See also Figure S3.

After 24 hours of differentiation (day 0 of the protocol), control cells lacked mCherry expression and formed compact colonies, whereas cells differentiated in the presence of Dox showed robust mCherry expression and adopted a more dispersed, mesenchymal-like morphology (Figure 3D). RT-qPCR confirmed strong induction of TBX6 expression (p<0.0001) and a concomitant reduction in *SOX2* levels (p = 0.0093; Figure S3A). TBX6 overexpression significantly upregulated markers associated with both paraxial mesoderm (*FOXC1*, *FOXC2*), and intermediate mesoderm (*OSR1*), suggesting activation of diverse mesodermal lineage programs (Figure S3A), consistent with features of an early, multipotent mesodermal state.

To define global transcriptional changes induced by acute TBX6 overexpression, we performed bulk RNA-seq after 24 hours of Dox treatment (Figures 3B,C and S3B,C). TBX6 induction resulted in widespread transcriptional rewiring, including downregulation of genes characteristic of pluripotency (*NANOG*), early mesendoderm (*EOMES*), and alternative lineage programs (endoderm regulator *FOXA2*), alongside upregulation of genes associated with differentiated mesodermal states (Figure 3E). These included somitic and paraxial mesoderm (*FOXC1, DLL1, ALDH1A2*), intermediate mesoderm (*OSR1, LHX1*) and cardiac/muscle lineages (*MYL7, NRAP*) (Figure 3E). Consistent with this, gene ontology (GO) analysis revealed enrichment of biological processes related to mesodermal differentiation and morphogenesis, including somite, renal nephron, and cardiac lineages (Figure S3D).

Gene set enrichment analysis (GSEA) using a reference human PCW2 embryonic atlas ^23^ demonstrated selective enrichment of differentiated mesodermal transcriptional programs following TBX6 induction, with highest enrichment for emergent, advanced and axial mesoderm signatures, alongside depletion of epiblast and primitive streak signatures (Figure 3E). In a PCW3 reference atlas ^24^, TBX6-induced programs aligned with multiple distinct mesodermal lineages, including paraxial and intermediate mesoderm, while neural programs were depleted (Figure S3E). Reciprocal analysis, in which a TBX6-induced gene module was mapped onto the PCW2 and PCW3 human embryonic atlases, further supported these findings (Figures S3F-M). Together, these data indicate that acute TBX6 overexpression is sufficient to promote transcriptional progression beyond early mesoderm toward differentiated mesodermal states, but does not enforce commitment to a single mesodermal lineage.

### Duration of TBX6 activity determines commitment to somitic or alternative mesodermal fates

Previous studies in mouse embryonic stem cells suggested that the duration of TBX6 expression can influence mesodermal differentiation ^39^. Since acute TBX6 induction activates transcriptional programs associated with multiple mesodermal lineages, we next tested whether altering the duration of TBX6 activity would influence mesodermal lineage commitment in hTLS. We first investigated the consequences of sustained TBX6 overexpression by differentiating PB-Dox-TBX6 cells to hTLS in the continuous presence of Dox (Figures 4A,B). As a control, iPSCs harbouring Dox-inducible mCherry alone were differentiated to hTLS (PB-Dox-mCherry; Figures 4A,B). At day 5, hTLS generated from both PB-Dox-TBX6 and PB-Dox-mCherry cells exhibited numerous mCherry^+^ cells, indicating sustained transgene expression. This suggests that the piggyBac-based Dox-inducible system remained unsilenced in these cells, likely due to the presence of multiple insulator sequences (Figures 4B-D).

**Figure 4.**
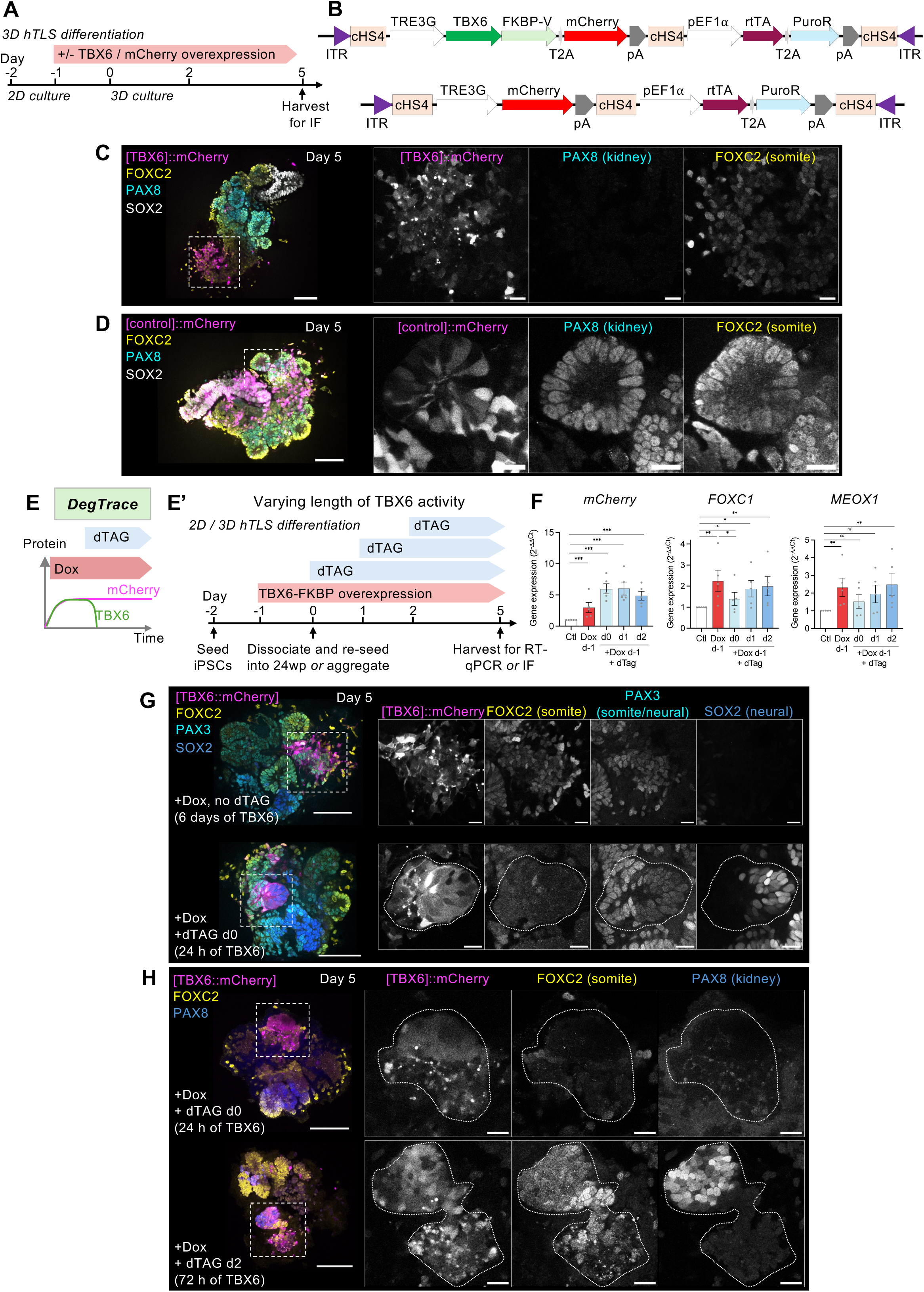
Duration of TBX6 activity determines commitment to a somitic mesoderm fate. (A) Experimental design to overexpress *TBX6* during the hTLS protocol in PB-Dox-TBX6 iPSCs. (B) Schematics of piggyBac-insertable constructs for Dox-inducible overexpression of TBX6-FKBP-V-T2A-mCherry (top), or mCherry alone (bottom). (C, D) Immunofluorescence of hTLS made with (C) PB-Dox-TBX6 iPSCs, or (D) PB-Dox-mCherry iPSCs, following sustained transgene induction (1 µM Dox from day -1), showing expression of SOX2 (grey), PAX8 (cyan), FOXC2 (yellow), and the mCherry transgene (magenta). Dotted white boxes indicate regions magnified in subsequent panels. Scale bars: main, 100 µm; insets, 20 µm. (E) Schematic of the DegTrace system. Dox induces TBX6-FKBP-T2A-mCherry expression, while subsequent dTAG treatment selectively depletes FKBP-fused TBX6 protein. mCherry fluorescence is retained, enabling lineage tracing of cells that transiently overexpressed TBX6. (E’) Experimental design to test various lengths of TBX6 protein activity in 2D- or 3D-cultured hTLS followed by RT-qPCR or IF analysis. (F) RT-qPCR for mCherry to assess transgene induction, or somitic mesoderm markers *FOXC1* and *MEOX1* in 2D-cultured hTLS generated from PB-Dox-TBX6 iPSCs either untreated (Ctl [control], white bars), treated with Dox only from day -1 (red bars), or treated with Dox from day -1 plus dTAG added either at day 0, day 1 or day 2 (blue bars). . Statistical significance was assessed using two-way ANOVA on log-transformed values, with treatment as the column factor and differentiation batch as the row factor; p value corresponds to the main effect of treatment (* p < 0.05, ** p < 0.01, *** p < 0.001, ns = not significant). Data are plotted as 2^-ΔΔCt^ for visualisation; bars represent mean ± SEM; dots represent mean of technical replicates from n = 5 independent differentiation batches per group. (G) Immunofluorescence of mCherry (magenta), FOXC2 (yellow), PAX3 (cyan) and SOX2 (blue) in representative day 5 PB-Dox-TBX6 hTLS. Top row: hTLS treated with Dox only from day -1 (6 days of TBX6 activity). Bottom row: hTLS treated with Dox from day -1 followed by dTAG from day 0 (24 h TBX6 activity). Dotted white boxes indicate regions of mCherry-expressing cells shown at higher magnification; an mCherry^+^ FOXC2^-^, PAX3^+^, SOX2^+^ epithelial structure is outlined in the bottom row. Scale bars: main, 100 µm; insets, 20 µm. (H) Immunofluorescence of mCherry (magenta), FOXC2 (yellow) and PAX8 (blue) expression in representative day 5 PB-Dox-TBX6 hTLS. Top row: hTLS treated with Dox from day -1 followed by dTAG from day 0 (24 h of TBX6 activity). Bottom row: hTLS treated with Dox from day -1 followed by dTAG from day 2 (72 h of TBX6 activity). Dotted white boxes indicate mCherry-expressing structures shown at higher magnification; mCherry^+^ epithelial structures are outlined with dotted lines in the insets. Scale bars: main, 100 µm; insets, 20 µm. See also Figure S4.

At day 5 in hTLS generated from PB-Dox-TBX6 cells, mCherry^+^ cells predominantly expressed the somitic marker FOXC2, lacked expression of PAX8 and SOX2, and adopted a mesenchymal morphology, indicating restriction toward a somitic mesoderm fate (Figure 4C). By contrast, hTLS generated from PB-Dox-mCherry cells exhibited mCherry expression within epithelial structures across somitic, renal, and neural compartments (Figure 4D), confirming that sustained TBX6 activity specifically, and not Dox-inducible transgene expression per se, drives a specific somitic lineage bias.

Since TBX6 activity is transient both *in vivo* and during hTLS development (Figures 2B and S2B) ^38^, we next tested whether restricting the duration of ectopic TBX6 activity would enable mesodermal diversification. To precisely control TBX6 protein stability while retaining a record of prior TBX6 expression, we utilised a FK506 binding protein (FKBP) domain fused to TBX6 to develop a conditional degradation and lineage-tracing system that we termed DegTrace (Figures 4E,E’) ^41^. In this system, Dox induces transcription of the TBX6-FKBP-T2A-mCherry transgene, whereas subsequent addition of dTAG selectively depletes TBX6 protein by engaging the fused FKBP domain with E3 ubiquitin ligases for proteasomal degradation ^41^, without affecting the co-translated mCherry reporter, thus enabling lineage tracing of cells that have transiently expressed TBX6 (Figures 4E,E’). We validated DegTrace in iPSCs, confirming that dTAG-mediated degradation selectively depleted TBX6 protein while preserving mCherry expression (Figures S4A,B).

We next applied DegTrace to 3D trunk differentiation. TBX6-FKBP-T2A-mCherry transgene overexpression was induced with Dox from day -1, and TBX6 protein degradation was triggered by dTAG addition at day 0 (after 24 h), day 1 (48 h), or day 2 (72 h) (Figures 4E,E’). RT-qPCR for the mCherry transcript showed that Dox alone induced transgene expression irrespective of subsequent dTAG addition (p < 0.001 versus control; Figure 4F). Sustained TBX6 overexpression (Dox alone) robustly induced the somitic mesoderm markers *FOXC1* and *MEOX1* (both p < 0.01 versus control; Figure 4F). By contrast, restricting TBX6 activity to the first 24 h led to diminished expression of these somitic markers, whereas inducing degradation after 48 or 72 h of TBX6 activity resulted in partial marker retention (Figure 4F).

Consistent with these transient transcriptional effects, immunofluorescence analysis revealed that cells exposed to transient TBX6 activity (24 h) failed to maintain mesodermal identity and instead adopted a SOX2^+^PAX3^+^ neural epithelial fate (Figures 4G,H and S4C,D), whereas cells exposed to an intermediate duration of TBX6 activity (72 h) differentiated into mesodermal epithelial structures expressing both the somitic mesoderm marker FOXC2 and the renal marker PAX8 (Figure 4H). By contrast, sustained TBX6 overexpression promoted mesenchymal differentiation and restriction to a FOXC2^+^PAX8^-^ somitic fate (Figures 4C,G).

Together, these findings indicate that a minimum duration of TBX6 activity is needed to suppress neural identity and promote mesodermal identity. An intermediate duration enables acquisition of a multipotent epithelial trunk mesoderm state with competence for intermediate (renal) mesoderm differentiation. By contrast, prolonged TBX6 activity biases cells toward a somitic mesoderm fate and reduces competence for alternative mesodermal lineages. This is accompanied by loss of epithelial characteristics, suggesting progression beyond an epithelial somite-like state toward a more mesenchymal somite derivative. Notably, the intermediate duration of exposure closely mirrors the endogenous dynamics of TBX6 expression observed in hTLS (Figures 2B and S2B,D).

### FOXC1 and FOXC2 establish and safeguard somitic mesoderm identity

Having established that TBX6 duration governs mesodermal fate output, we next sought to identify downstream mechanisms that stabilise specific mesodermal identities. TBX6 induction alone biased differentiation toward somite-like fates in hTLS, so we sought to investigate downstream transcriptional regulators of the somitic lineage. Analysis of the hTLS snRNA-seq mesodermal subset (Figure 1G) identified a somitic progenitor population enriched for canonical somitic transcription factors including *FOXC1*, *FOXC2*, *PAX3*, *TCF15* (Paraxis), *SIX1*, *MEOX1* and the Hedgehog pathway mediator *GLI2* (Figures 5A,B).

**Figure 5.**
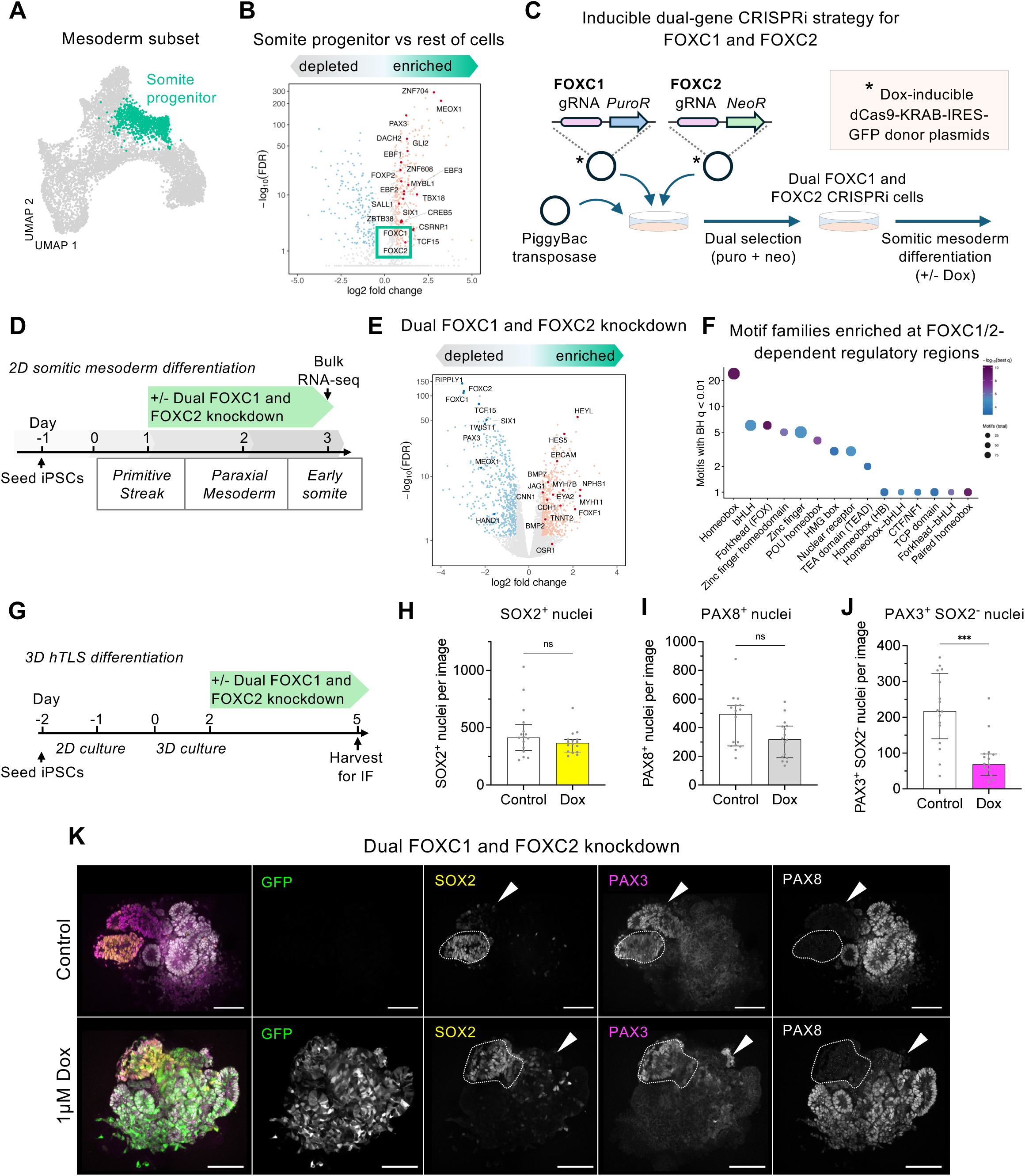
FOXC1 and FOXC2 establish and safeguard somitic mesoderm identity. (A) UMAP of 8,088 mesodermal subset cells, with the somite progenitor cluster (as in Figure 1G) highlighted. (B) Volcano plot of genes differentially expressed in the somite progenitor cluster compared with the remaining mesodermal subset. Genes with adjusted p-value < 0.05 and |log2 fold change| > 0.58 (±1.5-fold) are highlighted (red, upregulated; blue, downregulated). The top 20 upregulated transcription factors according to log2 fold change are shown in darker colour. Box highlights FOXC1 and FOXC2. (C) Inducible dual FOXC1 and FOXC2 CRISPRi strategy. Two piggyBac donor constructs each encoding a Dox-inducible dCas9-KRAB-IRES-GFP cassette and a single gRNA targeting either FOXC1 (with puromycin resistance; PuroR), or FOXC2 (with neomycin resistance; NeoR) were co-integrated into iPSCs using piggyBac transposase. Dual antibiotic selection yields a single cell line harbouring both cassettes. Dox treatment induces dCas9-KRAB expression, enabling simultaneous knockdown of *FOXC1* and *FOXC2* during somitic mesoderm differentiation. (D) Experimental design to test dual knockdown of *FOXC1* and *FOXC2* in a 2D somitic mesoderm differentiation protocol adapted from ^45^, followed by bulk RNA-seq. (E) Volcano plot of differential gene expression following dual knockdown of *FOXC1* and *FOXC2* in 2D somitic mesoderm differentiation (n = 3 independent differentiation batches per condition). Genes with adjusted p-value < 0.05 and |log2 fold change| > 0.58 (±1.5-fold) are highlighted (red, upregulated; blue, downregulated). Selected genes associated with somitic, kidney, and lateral plate/cardiomyocyte/smooth muscle identities are highlighted in darker colours. (Somitic: *FOXC1, FOXC2, MEOX1, HAND1, TCF15, TWIST1, RIPPLY1, SIX1, PAX3*; kidney: *OSR1, NPHS1, EYA2, EPCAM, CDH1, JAG1, HEYL, HES5*; lateral plate/cardiomyocyte/smooth muscle: *FOXF1*, *BMP2*, *BMP7*, *TNNT2*, *CNN1*, *MYH7B, MYH11*). (F) Dot plot showing enrichment of transcription factor motif families identified in regulatory regions of FOXC1/2-dependent genes versus expressed non-dependent genes (defined in (E)), using early somite ATAC-seq data ^45^. Dot position indicates the number of significantly enriched motifs per family (BH-adjusted q < 0.01); dot size reflects total motifs representation in HOMER; colour denotes the significance (-log_10_ q) of the most enriched motif in each family. (G) Experimental design to test dual knockdown of *FOXC1* and *FOXC2* in hTLS. (H-J) Quantification of the number of nuclei per confocal image that are (G) SOX2^+^, (H) PAX8^+^, or (I) PAX3^+^SOX2^-^, in FOXC1/2-dCas9-KRAB hTLS either untreated (control, white bars) or treated with 1µM Dox from day -1 (Dox, coloured bars). Dots represent n=16 individual hTLS across 2 independent batches per group; bars represent median ± IQR; Mann-Whitney test (*** p < 0.001). (K) Immunofluorescence of hTLS generated from FOXC1/2-dCas9-KRAB iPSCs and harvested at day 5, either untreated (control, top) or treated with Dox from day -1 (1 µM, bottom), showing GFP (green), SOX2 (yellow), PAX3 (magenta) and PAX8 (grey). Dotted white lines outline PAX3^+^SOX2^+^ neural structures, white arrowheads indicate PAX3^+^SOX2^-^ somitic tissue depleted upon Dox treatment. Scale bars, 100 µm. See also Figure S5.

We focussed on investigating the function of FOXC1 and FOXC2 in human embryoids based on prior genetic evidence that combined loss of these factors in mice disrupts somite formation and leads to expansion of intermediate (renal) mesoderm ^42^. As prior mouse studies indicated partial functional redundancy between FOXC1 and FOXC2 in early development ^42–44^, we engineered a dual knockdown system using dCas9-KRAB, hereafter referred to as FOXC1/2-dCas9-KRAB (Figures 5C and S5C).

To examine the role of FOXC1/2 specifically in somitic mesoderm differentiation, we adapted a published 2D human somitic mesoderm differentiation protocol ^45^ that recapitulates progression from primitive streak through paraxial mesoderm to an early somite-like state (Figures 5D and S5A,B). Dual knockdown of *FOXC1* and *FOXC2* was induced at the primitive streak stage, and cells were harvested at the early somite stage for bulk RNA-seq (Figures 5D and S5D,E).

Dual FOXC1/2 knockdown efficiently suppressed expression of both factors and was accompanied by a robust decrease in canonical somitic markers, including *RIPPLY1*, *PAX3*, *TCF15, HAND1* and *TWIST1* (Figure 5E). Concomitantly, dual FOXC1/2 knockdown induced genes associated with alternative mesodermal lineages. These included intermediate (renal) mesoderm (e.g. *NPHS1, EYA2, JAG1),* lateral plate mesoderm (*FOXF1, BMP2, BMP7*) and cardiomyocyte/smooth muscle-associated genes (e.g. *TNNT2*, *CNN1*, *MYH11, MYH7B*; Figures 5E and S5F). Primitive streak (*TBXT*, *CDX2*) and paraxial mesodermal genes (*TBX6, MSGN1*) were unchanged or upregulated, indicating that dual FOXC1/2 knockdown did not prevent establishment of paraxial mesoderm but instead impaired progression from paraxial mesoderm to an early somite state (Figure S5G). GO analysis confirmed depletion of patterning and skeletal development programs, with concomitant enrichment of contractile, heart, and renal system processes (Figure S5H).

To better understand how FOXC1/2-dependent genes correspond to human embryonic programs *in vivo*, we performed GSEA using PCW3-5 human embryonic reference atlases ^24^. This analysis revealed that FOXC1/2 knockdown led to depletion of signatures associated with somitic mesoderm derivatives and enrichment of alternative lineage programs, including intermediate (renal) mesoderm and cardiomyocyte signatures (Figure S5I). Moreover, GSEA using independent human embryonic heart ^46^ and kidney ^47^ reference atlases further demonstrated that genes upregulated upon FOXC1/2 knockdown were enriched for cardiomyocyte (Figure S5J) and renal nephron tube and podocyte signatures (Figure S5K). Together, these results indicate that FOXC1 and FOXC2 are required to establish somite transcriptional identity and to restrict alternative mesodermal differentiation programs.

To investigate mechanisms regulating FOXC1/2-dependent gene expression, we used published ATAC-seq profiles from the equivalent early somite population generated with the same protocol ^45^. Motif enrichment analysis was performed on accessible chromatin regions adjacent to genes we found to be downregulated following dual FOXC1/2 knockdown, using accessible regions associated with expressed genes that were not FOXC1/2-dependent as background. This analysis revealed enrichment of forkhead, homeobox, and basic helix-loop-helix (bHLH) motifs within FOXC1/2-dependent regulatory regions, suggesting that FOXC1 and FOXC2 act within a broader regulatory landscape of combinations of transcriptional regulators rather than as solitary lineage-determining factors (Figure 5F).

To test whether FOXC1/2 regulate somite identity in hTLS, we induced dual knockdown during hTLS differentiation (Figure 5G). While neural and renal tissues were preserved, PAX3^+^SOX2^-^ somitic tissue was markedly reduced (68% reduction compared to control; p < 0.0001), demonstrating a specific requirement for FOXC1/2 in establishing somitic mesoderm identity in hTLS (Figures 5H-K and S5L).

Finally, to determine whether TBX6 function relies on FOXC1/2 and thereby define their epistatic relationship, we generated iPSCs harbouring both a Dox-inducible TBX6 overexpression cassette and the FOXC1/2-dCas9-KRAB knockdown system (Figure S5M). TBX6 overexpression alone induced multiple somitic genes, including *TWIST1*, *TWIST2*, *MEOX1*, *MEOX2* and *TBX18*, however, this induction was abolished upon simultaneous FOXC1/2 knockdown, indicating that TBX6-driven somitic differentiation depends on FOXC1/2 activity (Figure S5N). By contrast, *MESP2,* previously described as a TBX6-responsive gene ^39^ was unaffected by FOXC1/2 knockdown, suggesting a parallel FOXC-independent TBX6 pathway (Figure S5N).

Together, these data indicate that mesodermal diversification is regulated by the duration of TBX6 activity, while FOXC transcription factors function downstream of TBX6 to constrain mesodermal plasticity and establish somitic identity.

### FOXC1 and FOXC2 stabilise somitic identity and preserve competence for further diversification

Having found that FOXC1/2 are required to establish somite mesoderm identity, we next asked whether they are additionally needed to maintain somite identity. To investigate this, we differentiated FOXC1/2-dCas9-KRAB cells to early somite stage and then induced dual knockdown (Figure 6A). Dual FOXC1/2 knockdown alone led to a reduction in somite markers *TCF15*, *TBX18* and *MEOX2,* whereas *MEOX1* and *SIX1* were unchanged, suggesting that FOXC1/2 loss partially destabilised the early somite program (Figure 6B).

**Figure 6.**
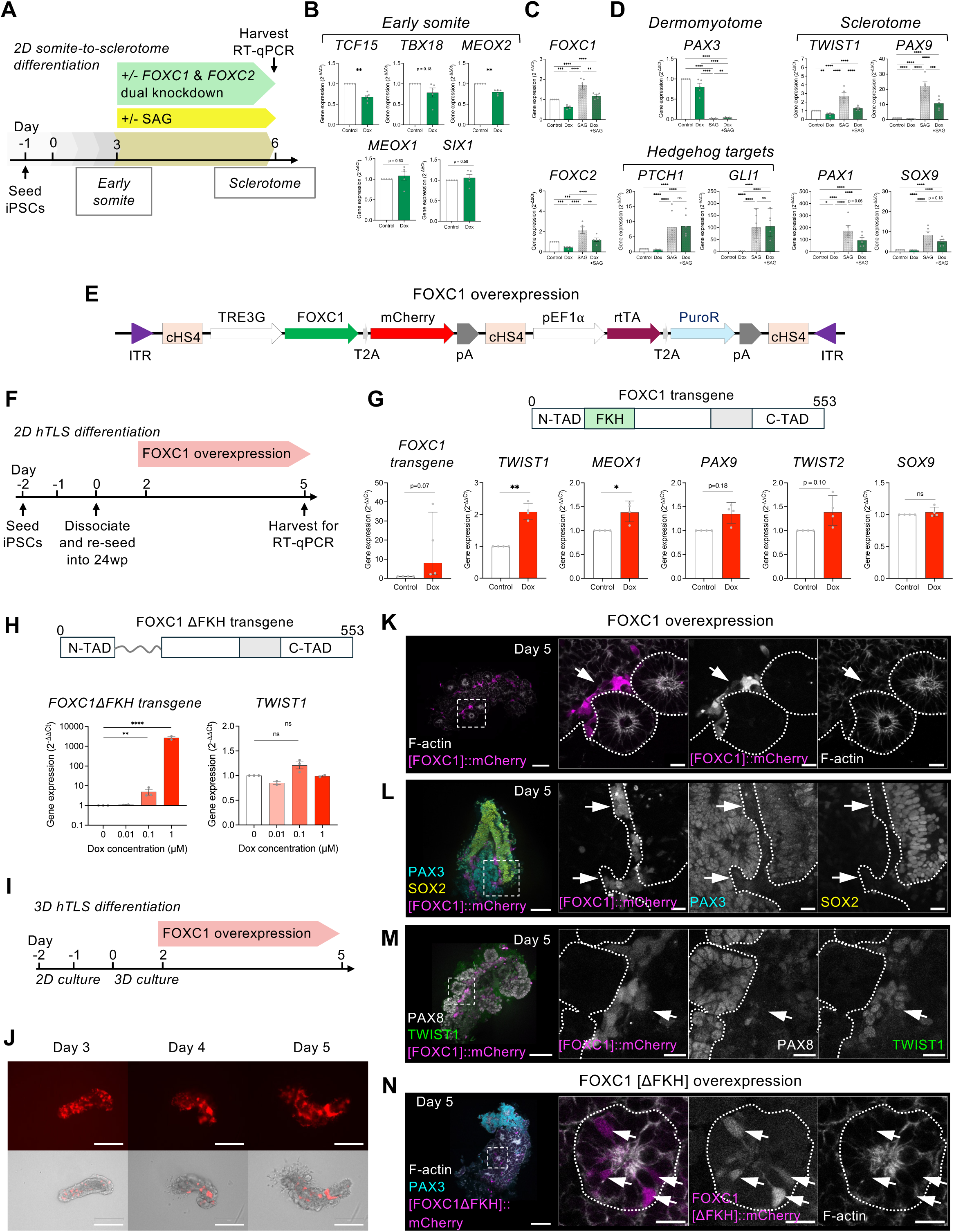
FOXC1 drives a partial sclerotome differentiation program. (A) Experimental design to test dual FOXC1 and FOXC2 knockdown in a 2D sclerotome differentiation protocol adapted from ^45^ followed by RT-qPCR. (B-D) RT-qPCR for B) early somite markers, C) *FOXC1* and *FOXC2*, and D) dermomyotome and sclerotome markers and canonical Hedgehog target genes, in FOXC1/FOXC2-dCas9-KRAB iPSCs differentiated to early somite and then either untreated (control) or treated from day 3 with doxycycline (Dox, B-D), or also with smoothened agonist (SAG), or combined Dox and SAG treatment (C,D). Statistical significance was assessed using two-way ANOVA on log-transformed values, with treatment as the column factor and differentiation batch as the row factor; p-values correspond to the main effect of treatment (* p < 0.05, ** p < 0.01, *** p < 0.001, **** p < 0.0001, ns = not significant). Data are plotted as 2^-ΔΔCt^; bars represent mean ± SEM; dots represent mean of technical replicates from n = 5 independent differentiation batches per group. (E) Schematic of the FOXC1 overexpression strategy (F) Schematic of experimental design to overexpress *FOXC1* in 2D-cultured hTLS for subsequent RT-qPCR. (G) Schematic of FOXC1 protein showing domain architecture, together with RT-qPCR analysis of *FOXC1* transgene, *TWIST1*, *MEOX1*, *PAX9, TWIST2* and *SOX9* expression in hTLS derived from PB-Dox-FOXC1 iPSCs at day 5, either untreated (control) or treated with Dox (0.1 µM) from day 2. Statistical significance was assessed using two-way ANOVA on log-transformed values, with treatment as the column factor and differentiation batch as the row factor; p-values shown correspond to the main effect of treatment (* p < 0.05, ** p < 0.01, *** p < 0.001, **** p < 0.0001). Data are plotted as 2^-ΔΔCt^ for visualisation; bars represent mean ± SEM; dots represent mean of technical replicates from n = 4 independent differentiation batches per group. N-TAD, N-terminal activation domain; FKH, forkhead domain; C-TAD, C-terminal activation domain; grey box, putative inhibitory domain ^73^. (H) Schematic of FOXC1[ΔFKH] protein showing domain architecture with FKH DNA-binding domain replaced with a flexible linker; plots show RT-qPCR analysis for transgene and TWIST1 in day 5 2D-hTLS generated from iPSCs harbouring Dox-inducible FOXC1[ΔFKH] and treated with varying concentrations of Dox (0, 0.01, 0.1, 1 µM) from day 2. Statistical significance was assessed using two-way ANOVA on log-transformed values (ΔCt), with treatment as the column factor and differentiation batch as the row factor. The p values shown correspond to the main effect of treatment (** p < 0.01, **** p < 0.0001). Data are plotted as 2^-ΔΔCt^; bars represent mean ± SEM; dots represent mean of technical replicates from n=2-3 independent differentiation batches per group.N-TAD, N-terminal activation domain; FKH, forkhead domain; C-TAD, C-terminal activation domain; grey box, putative inhibitory domain ^73^. (I) Experimental design to overexpress FOXC1 in hTLS (J) Brightfield and corresponding red fluorescence images of hTLS generated from PB-Dox-FOXC1 iPSCs at days 3, 4, and 5. Scale bars, 300 µm. (K) Immunofluorescence of hTLS generated from PB-Dox-FOXC1 iPSCs following FOXC1 overexpression from day 2, stained for mCherry (magenta) and F-actin (phalloidin, grey). White dotted box indicates region shown at higher magnification, in which individual epithelial cysts are outlined with dotted lines. Arrows indicate FOXC1-overexpressing cells occupying a subepithelial position. Scale bars: main, 100 µm; insets, 20 µm. (L) Immunofluorescence of hTLS generated from PB-Dox-FOXC1 iPSCs following FOXC1 overexpression from day 2, stained for mCherry (magenta), PAX3 (cyan), and SOX2 (yellow). Arrows indicate FOXC1-overexpressing cells lacking both PAX3 and SOX2. Scale bars: main, 100 µm; insets, 20 µm. (M) Immunofluorescence of hTLS generated from PB-Dox-FOXC1 iPSCs following FOXC1 overexpression from day 2, stained for mCherry (magenta), PAX8 (grey), and TWIST1 (green). Arrows indicate FOXC1-overexpressing cells co-expressing TWIST1. Scale bars: main, 100 µm; insets, 20 µm. (N) Immunofluorescence of hTLS generated from PB-Dox-FOXC1[ΔFKH] iPSCs following overexpression from day 2, stained for mCherry (magenta), PAX3 (cyan), and F-actin (phalloidin, grey). White dotted box indicates region shown at higher magnification, in which individual epithelial cysts are outlined with dotted lines. Arrows indicate FOXC1[ΔFKH]-overexpressing cells within epithelial cysts. Scale bars: main, 100 µm; insets, 20 µm. See also Figures S6 and S7.

We next tested whether dual knockdown affected the ability of early somite cultures to differentiate into sclerotome, a transient embryonic structure that gives rise to the vertebral skeleton. Studies of avian and mouse development have established that sclerotome differentiation *in vivo* is triggered in the ventral somite by Hedgehog ligands derived from the notochord and floorplate ^48,49^. This process is characterised by repression of PAX3, upregulation of FOXC1/2, and induction of downstream markers including TWIST1, PAX1, PAX9, and SOX9 together with an epithelial-to-mesenchymal transition (EMT) (Figure S6A).

We adapted a 2D human somite-to-sclerotome differentiation protocol in which early somite cultures were exposed to smoothened agonist (SAG) to activate Hedgehog signalling and trigger sclerotome differentiation (Figure 6A) ^45^. SAG treatment resulted in efficient generation of sclerotome characterised by suppression of *PAX3* and upregulation of *FOXC1*, *FOXC2*, *TWIST1*, *PAX9*, *PAX1* and *SOX9* compared to untreated cells (Figures 6A,C,D).

Dual FOXC1/2 knockdown induced at the early somite stage, in combination with SAG treatment, strongly attenuated induction of sclerotome markers (Figures 6C,D), whereas induction of canonical Hedgehog pathway targets *PTCH1* and *GLI1* by SAG was unaffected by FOXC1/2 knockdown (Figure 6D). These data indicate that FOXC1/2 are not necessary for Hedgehog signalling per se, but instead support maintenance of somite identity and thereby preserve competence for sclerotome differentiation, or indeed may play a more direct role in sclerotome differentiation.

As FOXC1/2 knockdown inhibited the establishment and maintenance of somite identity, we next asked whether they are sufficient to promote somite differentiation when overexpressed in hTLS. Given the functional redundancy of FOXC1 and FOXC2 *in vivo* and in hTLS, we focussed on FOXC1 and generated a Dox-inducible FOXC1 overexpression line (PB-Dox-FOXC1; Figure 6E). FOXC1 overexpression was induced at day 2 of differentiation to match the timeframe of the previous FOXC1/2 knockdown experiments in hTLS (Figures 5G-K and 6F).

RT-qPCR analysis of 2D-cultured hTLS generated with PB-Dox-FOXC1 cells revealed that FOXC1 overexpression robustly upregulated *TWIST1* (∼2-fold; p < 0.01), a bHLH transcription factor associated with ventral somite identity, sclerotome differentiation and EMT (Figure 6G). FOXC1 overexpression also upregulated the ventral somite marker *MEOX1* (∼1.4-fold; p < 0.05), but only modestly induced *PAX9* and *TWIST2* and failed to induce *SOX9* (Figure 6G).

Overexpression of a FOXC1 variant lacking the forkhead DNA-binding domain (FOXC1ΔFKH) failed to upregulate *TWIST1,* even at the highest Dox concentration (1 µM) despite robust transgene induction, indicating that *TWIST1* induction is specific to transcriptionally active FOXC1 (Figure 6H).

To investigate the effects of FOXC1 overexpression in a context more closely resembling native embryonic tissue, PB-Dox-FOXC1 cells were differentiated to 3D-hTLS, and FOXC1 overexpression was induced from day 2 of hTLS differentiation (Figure 6I). Robust, heterogeneous mCherry induction was observed in elongating hTLS and maintained through days 3-5 (Figure 6J). FOXC1-overexpressing cells were largely excluded from the epithelial cysts that formed and instead occupied a sub-epithelial position, consistent with acquisition of a mesenchymal-like state (Figure 6K). By contrast, FOXC1ΔFKH-overexpressing cells were observed within epithelial cysts, indicating that FOXC1 transcriptional activity is needed to drive the mesenchymal phenotype (Figure 6N).

Cells overexpressing wild type FOXC1 (FOXC1::mCherry^+^) within hTLS did not express SOX2, PAX3, or PAX8, indicating that FOXC1 overexpression does not promote neural, dorsal somite, or renal fates (Figures 6L,M). Moreover, FOXC1-overexpressing cells robustly expressed TWIST1 (Figure 6M) but rarely co-expressed the sclerotome marker SOX9 (Figure S6H), suggesting that FOXC1 is insufficient to fully execute the sclerotome differentiation program. Overexpression of FOXC2 yielded a similar phenotype, with sub-epithelial localisation of overexpressing cells and absence of SOX2 and PAX8 expression (Figure S6I).

Consistent with this, snRNA-seq analysis of untreated hTLS did not reveal a sclerotome population (Figures S1G-J), despite expression of the Hedgehog pathway mediators *GLI2* and *GLI3* (Figures S6A,B). This likely reflects the absence of notochord-and floorplate-derived Hedgehog ligands in this system (Figures S6A,B). To test whether hTLS retain competence for sclerotome differentiation, we activated Hedgehog signalling with SAG from day 5 to day 8 of differentiation (Figure S6C). SAG treatment led to expansion of FOXC2^+^ tissue and depletion of PAX3^+^ dorsal somitic domains, consistent with somite ventralisation and sclerotome differentiation (Figures S6D-G).

Together, these results indicate that FOXC1 and FOXC2 are required to establish and stabilise somitic mesoderm identity and to enable somite-to-sclerotome differentiation. Moreover, while FOXC1 overexpression is sufficient for *TWIST1* induction and EMT-like behaviour, full sclerotome differentiation requires additional cues such as Hedgehog signalling.

### TWIST1 is required for Hedgehog-driven sclerotome differentiation

Because our ATAC-seq analysis suggested that FOXC factors co-operate with bHLH factors to establish the somitic transcriptional landscape, we hypothesised that a bHLH factor might act as the direct effector of FOXC1/2. FOXC1 overexpression upregulated the bHLH factor TWIST1 and drove EMT in hTLS, and TWIST1 has known roles in inducing EMT ^50,51^, positioning TWIST1 as a plausible candidate bHLH functioning downstream of FOXC1. We edited iPSCs to harbour a Dox-inducible TWIST1 overexpression cassette and differentiated them to hTLS (Figures S7A,B). Inducible TWIST1 overexpression led to sub-epithelial localisation of TWIST1::mCherry^+^ cells and exclusion from epithelial cysts, closely resembling the FOXC1 overexpression phenotype (Figure S7C), suggesting that TWIST1 may function downstream of FOXC1 to drive EMT in this system.

Finally, as FOXC1 and FOXC2 are necessary for sclerotome differentiation, we asked whether TWIST factors are also required. Dual knockdown of TWIST1 and TWIST2 during SAG-induced differentiation (Figure S7D) strongly reduced SAG-induced expression of sclerotome markers, but did not affect induction of *FOXC1* or *FOXC2* by SAG (Figure S7E). Moreover, SAG-induced expression of *PTCH1* and *GLI1* was unaffected by TWIST1/2 knockdown (Figure S7E). Together, these data suggest that FOXC factors may promote EMT via TWIST factors, and support a model in which TWIST factors do not affect Hedgehog signalling per se, but instead act as necessary effectors of Hedgehog-driven sclerotome differentiation in human embryonic mesoderm.

## Discussion

How lineage identities first emerge and diversify during early human embryogenesis remains a fundamental unresolved question, largely owing to the experimental inaccessibility of post-gastrulation human development *in vivo*. While recent single-cell atlases have mapped transcriptomic states during early human development, experimental dissection of the mechanisms governing lineage diversification during this window has remained limited. To address this knowledge gap, we use an iPSC-derived hTLS embryoid model, combined with genetic tools that enable precise temporal control of transcription factor activity with dual gene knockdown, to uncover a hierarchical regulatory logic underlying early human mesodermal diversification. Our data support a model in which TBX6 functions as a “temporal gate” governing mesodermal diversification, with the duration of TBX6 activity determining intermediate (renal) versus somitic mesodermal fates (Figure 7). Downstream of TBX6, FOXC1/2 act as a “lock” to establish and stabilise somitic mesoderm identity, preserving the ability of somite cells to undergo sclerotome differentiation in response to Hedgehog signalling, in part through induction of TWIST factors. Given that many developmental transcription factors are transiently expressed, our findings suggest that the duration of transcription factor activity may act as a temporal parameter that helps define competence windows to hierarchically restrict lineage plasticity during embryogenesis.

**Figure 7.**
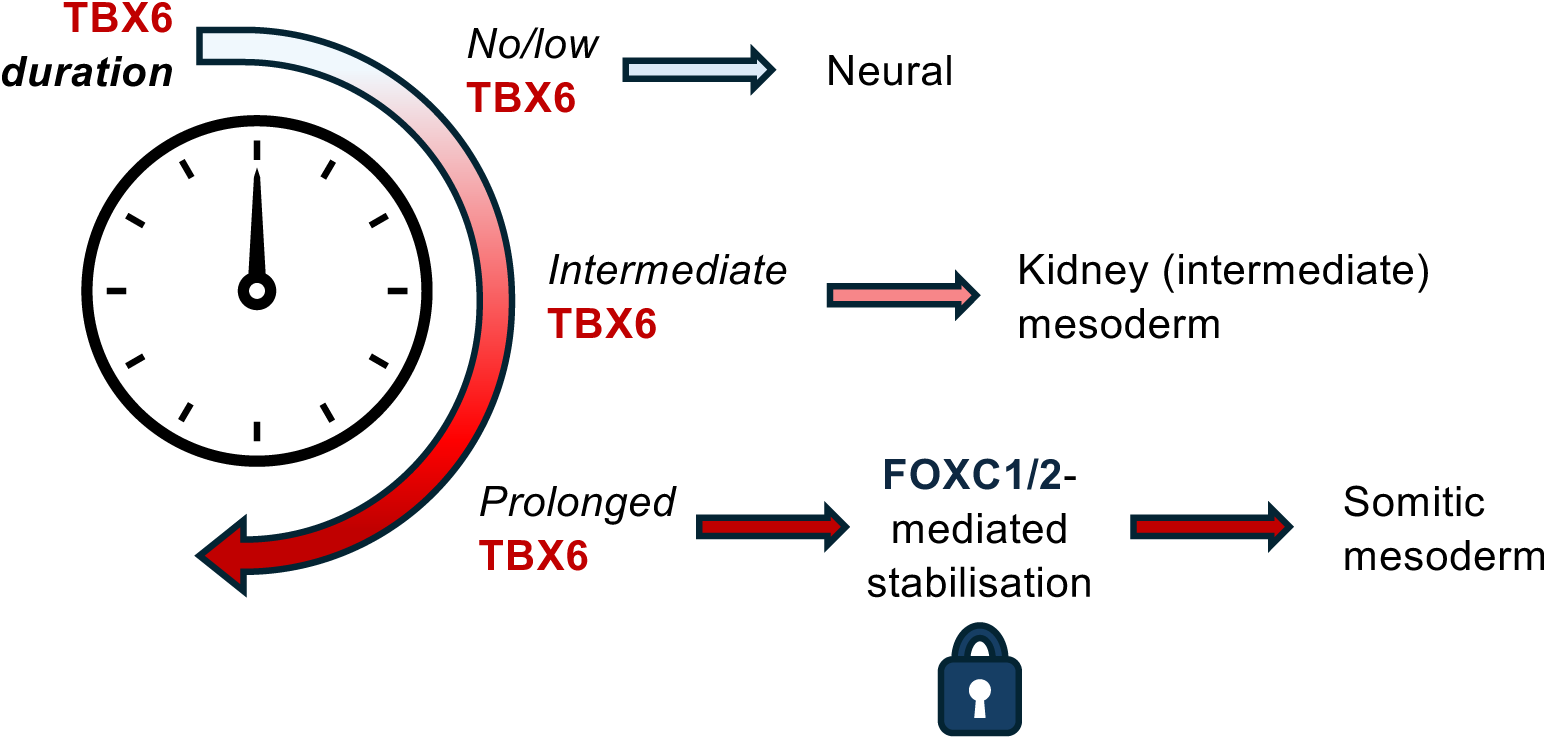
Proposed temporal gate and lock mechanism for human mesodermal diversification. In this model, TBX6 activity duration functions as a temporal gate that determines lineage competence during early human mesoderm specification. In the absence or at low levels of TBX6, cells adopt a neural fate, whereas intermediate TBX6 activity permits differentiation into intermediate (renal) mesoderm. Prolonged TBX6 activity biases cells toward a somitic mesodermal fate. Downstream, FOXC1/2 act as a stabilising “lock” that consolidates somitic identity, restricting alternative lineage potential and preserving competence for subsequent differentiation. Together, these findings define a hierarchical regulatory logic in which TBX6 activity duration regulates competence for mesodermal diversification, followed by downstream stabilisation of committed cell identities.

We benchmarked our embryoid model against human *in vivo* embryonic reference atlases, validating hTLS as an experimentally tractable model that recapitulates key aspects of post-gastrulation human embryonic development. The presence of paraxial (somitic) and intermediate (renal) mesoderm in our hTLS model enabled investigation of the logic underpinning the emergence of these distinct mesodermal subtypes. Fate-mapping of peri-gastrulation avian and mouse embryos showed that mesodermal subtypes including paraxial and intermediate mesoderm arise from discrete positions along the rostral-caudal axis of the primitive streak ^7–10,13^. Moreover, rostrally-located streak cells fated to become head mesenchyme and anterior somites would adopt lateral fates including intermediate and lateral plate mesoderm when grafted caudally ^7^, indicating developmental equivalence within a multipotent cellular pool. While morphogen gradients including BMP and FGF pattern the mediolateral axis of the mesoderm ^3,52^, specific transcription factors that stabilise lineage identity remain poorly defined. Our finding that FOXC1/2 promotes commitment to somitic mesoderm and restricts alternative fates including intermediate mesoderm provides a molecular basis for mesodermal fate commitment likely operating downstream of local microenvironmental cues. Our epistasis experiments, and our finding that *FOXC1* and *FOXC2* are induced by acute TBX6 overexpression, position FOXC1/2 functionally downstream of TBX6, either through direct chromatin-level regulation or via intermediate factors.

Our DegTrace system enabled precise control over the duration of transcription factor activity while simultaneously tracing the fate of modified cells, allowing us to experimentally interrogate how transcription factor dynamics influence lineage outcomes. Using this system, we uncovered a duration-dependent TBX6 regulatory logic underlying mesodermal diversification. TBX6 is classically described as a critical regulator of somitic mesoderm specification. TBX6 null mice lack somitic mesoderm and instead exhibit three neural tubes ^53,54^, and TBX6 has been proposed to repress the *SOX2* locus to permit somitic mesoderm differentiation in NMPs ^54^. Accordingly, recent human population genetics studies have revealed *TBX6* variants to be associated with congenital vertebral malformations, reflecting the somitic origin of vertebrae ^55–57^. Our finding that FOXC1/2 act downstream of TBX6 to establish and stabilise somitic mesoderm identity provides a mechanistic explanation for these defects. Consistent with this, *FOXC1* mutations in humans can cause Axenfeld-Rieger syndrome, which includes skeletal abnormalities such as scoliosis in addition to ocular defects ^58,59^.

Our observation that *TBX6* knockdown depletes both kidney and somite mesoderm is consistent with a model in which a multipotent mesodermal population expressing *TBX6* gives rise to both intermediate and paraxial mesoderm during post-gastrulation development. Studies of early mouse development mapped initial *Tbx6* expression broadly throughout the primitive streak, becoming progressively more posteriorly restricted as development proceeds ^38^. A multipotent TBX6^+^ progenitor of intermediate and somitic mesoderm is supported by lineage tracing studies of *Tbx6*-expressing cells in mice, which showed labelling of somitic and intermediate mesoderm when traced at early stages, and labelling restricted to somites if traced later ^60^. These observations support a model in which longer durations of TBX6 activity bias cells toward somitic fate, whereas shorter exposure permits alternative mesodermal outcomes. Supporting this broader role, *TBX6* variants have been associated with defects in intermediate mesoderm derivatives including the kidneys, in both human genetic studies and mouse models ^61–66^. Together with these studies, our findings position TBX6 as a potential regulator of intermediate mesoderm development in addition to its known role as a somitic mesoderm specifier.

The duration-dependent effects of TBX6 activity revealed by DegTrace are consistent with a recent report showing that a defined pulse of TBX6 induction enhances cardiomyocyte differentiation efficiency in pluripotent stem cell protocols ^39^. These observations raise the question of how the activity duration of a single transcription factor can encode competence for divergent lineage outcomes. One possibility is that prolonged TBX6 exposure permits accumulation of downstream cofactors that modify transcriptional output, whereas transient exposure fails to stabilise these networks. Additionally, mutually repressive interactions may occur between downstream targets as has recently been suggested for *Osr1* and *Foxc2* in mouse ^66^. Dissecting the chromatin and network dynamics underlying this temporal encoding will be an important direction for future work. The relative resistance of our piggyBac system to transgene silencing in many cells enabled inducible, sustained TBX6 activity, highlighting its utility for interrogating temporal transcription factor dynamics in differentiating systems, and offers advantages over recently described non-inducible transgene expression systems ^67^.

Our findings support a model in which FOXC1 and FOXC2 establish and stabilise somite identity in part by restricting alternative lineage programs. Comparison with human embryonic reference atlases revealed that dual FOXC1/2 knockdown caused loss of canonical somite markers alongside induction of intermediate mesoderm and cardiomyocyte/smooth muscle-associated gene programs. This is consistent with a shift toward alternative mesodermal fates when somite development is impaired, suggesting that FOXC1/2 help restrict, and may actively repress, intermediate and lateral plate mesoderm identities during development. In support of this, dual FOXC1/2 knockdown upregulated *FOXF1,* a lateral plate mesoderm marker *in vivo* ^68^, and *BMP2* and *BMP7,* which regulate lateral plate mesoderm development ^69^. Our findings contrast with prior studies of *Foxc1/2* double knockout mice which exhibited loss of both somitic and cardiac lineages ^44^, suggesting potential divergence in FOXC1/2 function between human and mice and highlighting the importance of using human-specific models, including hTLS, to study early human developmental mechanisms.

Notably, FOXC1 overexpression alone was not sufficient to drive somite differentiation. Our analysis of published ATAC data of early somite cultures revealed that FOXC motifs were co-enriched with bHLH and homeobox factors specifically at FOXC1/2-dependent genes in open chromatin compared to non-dependent genes. It is therefore likely that FOXC1 acts in a broader combinatorial regulatory framework to establish somite identity that may require cooperativity with bHLH transcription factors. We found that potential FOXC1/2 targets included bHLH factors such as *TCF15* and *HAND1* and homeobox factors including *MEOX1*, which could function as cooperative partners with FOXC1/2 in somite differentiation. Moreover, FOXC1 overexpression drove induction of the bHLH factor *TWIST1* and EMT-like behaviour. Thus, FOXC1-TWIST1 cooperativity may have a specific role in regulating the acquisition of a migratory, mesenchymal phenotype during sclerotome differentiation. However, *FOXC1* expression is not sufficient to drive full sclerotome differentiation, and additional cues such as Hedgehog signalling are needed. The established role of FOXC1 in cancer-associated EMT ^70,71^ suggests that this developmental function in promoting mesenchymal transition may be co-opted in pathological contexts.

In summary, our study reveals a duration-dependent hierarchical regulatory logic governing human mesodermal diversification and highlights the utility of iPSC-derived embryoid models, combined with transgene overexpression and genetic perturbations, for gaining insights into otherwise inaccessible stages of early human embryogenesis.

---

**Supplementary Figure 1.**
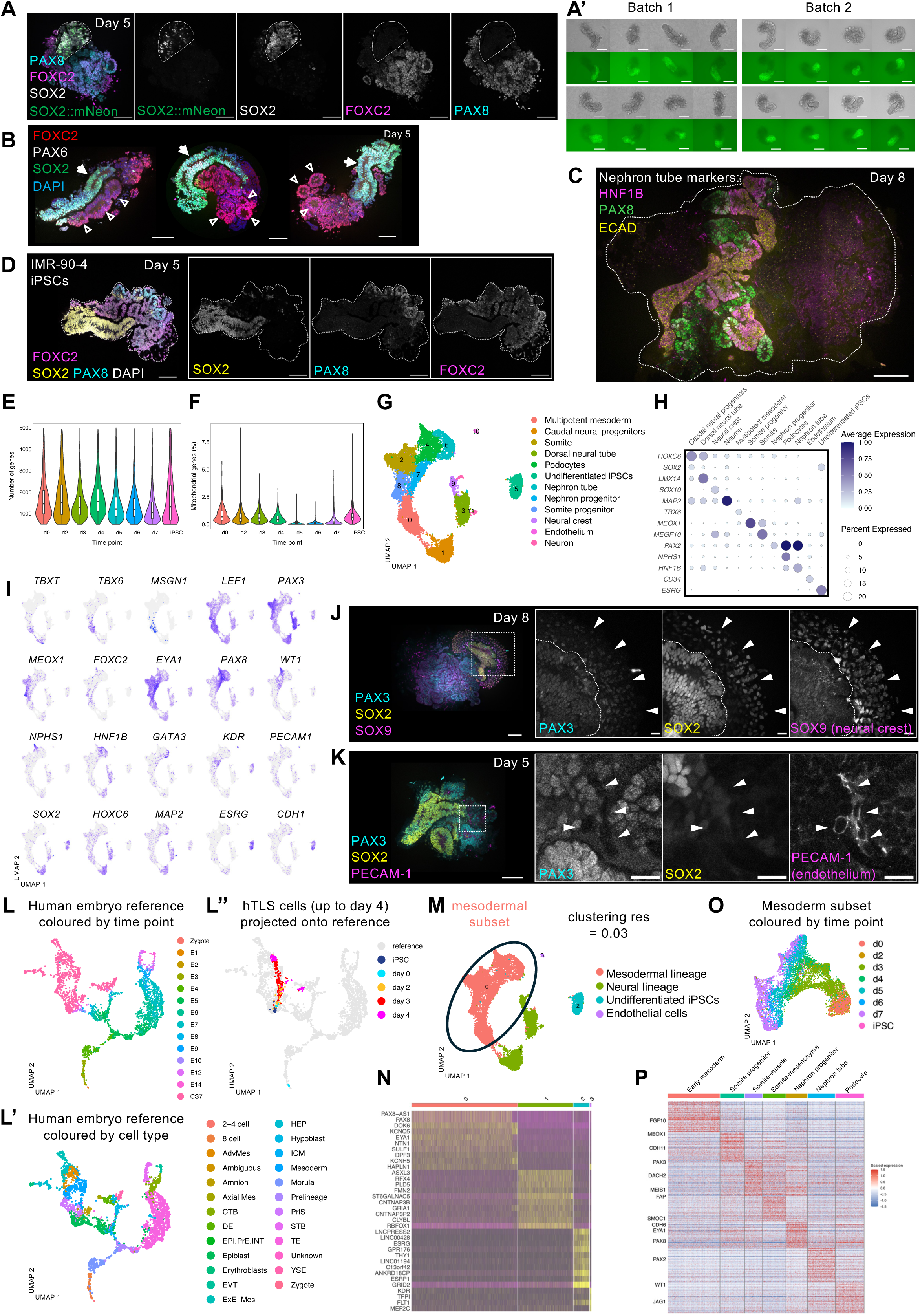
snRNA-seq quality control and additional immunofluorescence analyses (related to Figure 1) (A) Immunofluorescence of day 5 hTLS generated from SOX2::mNeon reporter iPSCs showing endogenous mNeon signal (green), and staining for SOX2 (grey), FOXC2 (magenta) and PAX8 (cyan) expression. A SOX2^+^ neural tube-like structure is outlined with a dotted line. mNeon expression is specific to SOX2^+^ cells within the neural tube-like structure but does not label all SOX2^+^ cells. Scale bars, 100 µm. (A’) Representative hTLS from two independent differentiation batches generated using SOX2::mNeon iPSCs, shown in brightfield and corresponding green fluorescence to illustrate intra- and inter-batch variability. (B) Immunofluorescence of FOXC2 (red), PAX6 (white) and SOX2 (green) expression in hTLS at day 5, with nuclei counterstained with DAPI (blue). Arrows indicate SOX2^+^ PAX6^+^ neural tube-like structures; arrowheads indicate FOXC2^+^ epithelial somite-like structures. Scale bars, 100 µm. (C) Immunofluorescence of the nephron markers PAX8 (green), ECAD/CDH1 (yellow), and HNF1B (magenta) in a day 8 hTLS. hTLS is outlined with dotted white line. Scale bar, 100 µm. (D) Immunofluorescence of PAX8 (cyan), SOX2 (yellow), and FOXC2 (magenta) in a day 5 hTLS derived from an independent iPSC line, IMR-90-4. Nuclei are counterstained with DAPI, grey. hTLS is outlined with dotted line. Scale bars, 100 µm. (E) Violin plots of hTLS snRNA-seq data showing the number of detected genes per cell at each time point. (F) Violin plots of hTLS snRNA-seq data showing the percentage of mitochondrial reads per cell at each time point. (G) UMAP representation of 13,618 hTLS cells coloured by unsupervised clustering with annotations determined by marker gene expression. (H) Dot plot showing expression of selected marker genes for each cluster in (G). (I) UMAP plots of 13,618 hTLS cells as in (G), with expression of selected marker genes overlaid. (J) Immunofluorescence of PAX3 (cyan), SOX2 (yellow), and the neural crest marker SOX9 (magenta) in hTLS at day 8. White box indicates magnified region. Dotted white line in the insets outline a PAX3^+^SOX2^+^ neural tube-like structure; arrowheads indicate delaminating SOX9^+^ neural crest cells. Scale bars: main image, 100 µm; insets, 20 µm. (K) Immunofluorescence of the neural/somite marker PAX3 (cyan), the neural marker SOX2 (yellow) and the endothelial marker PECAM-1/CD31 (magenta), in hTLS at day 5. White box indicates magnified region. Arrowheads indicate a PECAM-1^+^ vessel-like structure nestled between SOX2^+^ and PAX3^+^ neural/somitic tissue. Scale bars: main image, 100 µm; insets, 20 µm. (L, L’, L’’) Transcriptional similarity between hTLS snRNA-seq dataset and human reference embryogenesis dataset from Zhao et al., ^25^. UMAP representations of human reference embryogenesis dataset coloured by (L) time point and (L’) cell type using annotations from the reference. (L’’) projection of hTLS snRNA-seq cells (iPSC and days 0-4) onto reference UMAP, coloured by time point, with reference cells in grey. (M) UMAP of hTLS cells as in (G), coloured by unsupervised clustering at resolution = 0.03 to identify mesodermal cells (cluster 0), which were subsequently sub-clustered to isolate the mesodermal subset. (N) Heatmap of top 10 marker genes defining clusters in (M). Enriched genes were first determined with Wilcoxon rank sum test with thresholds log2 fold change > 1 and adjusted p-value < 1e-8. (O) UMAP of mesodermal subset (as in Figure 1G) coloured by time point. (P) Heatmap showing top 50 genes defining each cluster of the mesodermal subset annotated as in Figure 1G, with selected marker genes presented. Enriched genes were first determined with Wilcoxon rank sum test with thresholds log2 fold change > 1 and adjusted p-value < 0.005.

**Supplementary Figure 2.**
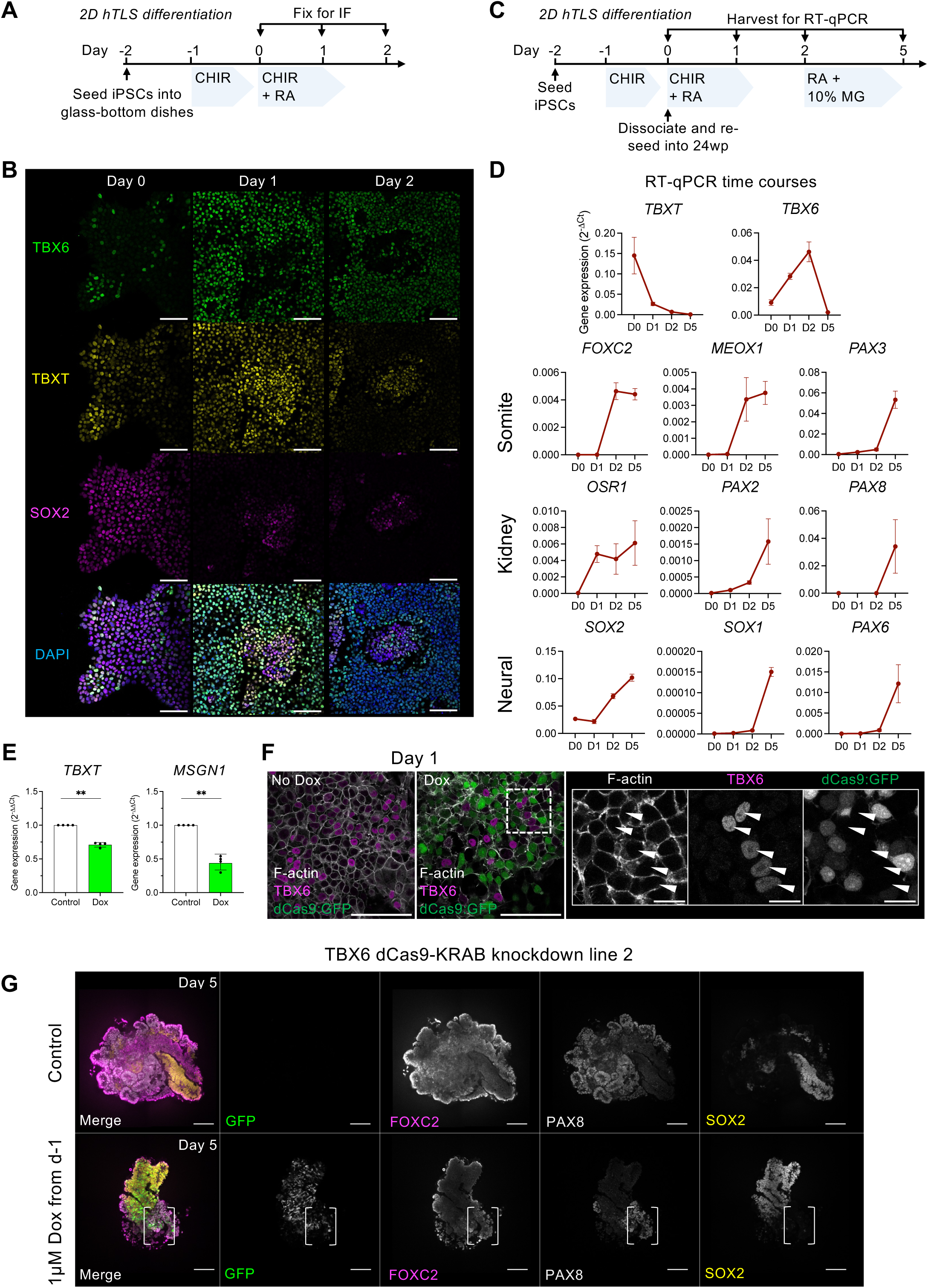
TBX6 is expressed transiently throughout the mesoderm before mesoderm diversification (related to Figure 2) (A) Experimental design to differentiate iPSCs in the hTLS protocol on glass-bottomed dishes without aggregation for subsequent immunofluorescence analysis. (B) Immunofluorescence of differentiated iPSCs (as outlined in A) showing expression of TBX6 (green), TBXT/Brachyury (yellow) and SOX2 (magenta) expression, with nuclei counterstained with DAPI (blue). Scale bars, 100 µm. (C) Strategy to differentiate iPSCs using the hTLS protocol followed by dissociation then reseeding at day 0 to generate hTLS pools maintained in 2D culture for subsequent RT-qPCR analysis. (D) Time-course RT-qPCR analysis of selected of early mesoderm, somite, kidney and neural markers during the hTLS protocol. Dots and bars represent mean ± SEM; n = 4 wells of hTLS per time point across 2 independent differentiation batches. (E) RT-qPCR analysis of *TBXT* and *MSGN1* expression in TBX6-dCas9-KRAB iPSCs differentiated using the hTLS protocol and harvested after 24 hours (as in Figure 2E), either untreated (control) or treated with Dox (1 µM) from day -1. Dots represent n = 4 independent differentiation batches. Two-way ANOVA on log-transformed values (ΔCt), with treatment as the column factor and differentiation batch as the row factor. P values correspond to treatment effect (** p < 0.01). Data are plotted as 2^-ΔΔCt^; bars represent mean ± SEM. (F) Immunofluorescence of TBX6-dCas9-KRAB iPSC cells differentiated as in (A) either without or with Dox (1 µM) treatment from day -1 to induce TBX6 knockdown, showing expression of TBX6 (magenta), and F-actin (grey) to outline cells, with GFP (green) as a readout for expression of the knockdown cassette, which is expressed heterogeneously after Dox treatment. Dotted box indicates magnified region. Arrowheads indicate expression of TBX6 in cells that lack GFP expression and thus did not express the knockdown cassette. Scale bars main images 100 µm; insets, 20 µm. (G) Immunofluorescence of day 5 hTLS generated from a second independent TBX6-dCas9-KRAB iPSC line with a distinct gRNA to target TBX6, either with no treatment (control, top) or following Dox treatment from day -1 (1µM Dox, bottom), showing GFP (green), SOX2 (yellow), FOXC2 (magenta), and PAX8 (grey). White brackets indicate FOXC2^+^PAX8^+^ cells confined to a GFP^-^ region. Scale bars, 100µm.

**Supplementary Figure 3.**
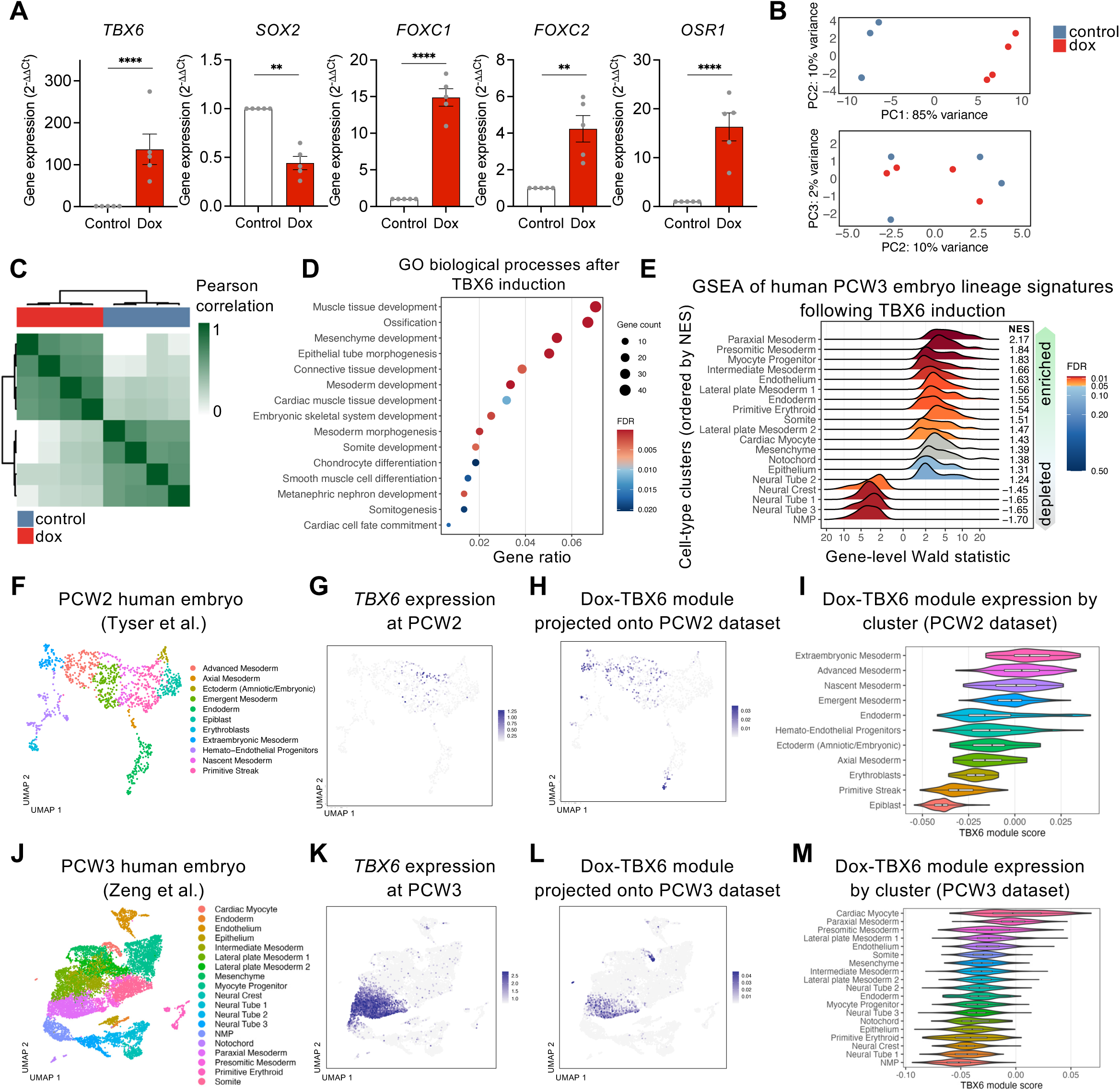
TBX6 RNA-seq quality control and additional analysis (related to Figure 3) (A) RT-qPCR analysis of *TBX6, SOX2,* somitic mesoderm markers *FOXC1* and *FOXC2,* and the intermediate mesoderm marker *OSR1,* in PB-Dox-TBX6 iPSCs differentiated using the hTLS protocol and harvested after 24 h, either untreated (control) or treated with Dox (0.1 µM) from day -1. Dots represent n = 5 independent differentiation batches. Two-way ANOVA on log-transformed values (ΔCt), with treatment as the column factor and differentiation batch as the row factor. P values correspond to treatment effect (** p < 0.01, **** p < 0.0001). Data are plotted as 2^-ΔΔCt^; bars represent mean ± SEM. (B) Principal component plots of the TBX6 overexpression bulk RNA-seq dataset. (C) Sample-to-sample correlation heatmap of the TBX6 overexpression bulk RNA-seq dataset. (D) Dot plot showing gene ontology (GO) biological processes (BP) analysis, highlighting selected processes upregulated following TBX6 overexpression. Dot size indicates gene count; colour denotes FDR-adjusted p-value. (E) Ridge plots of gene set enrichment analysis (GSEA) of genes following TBX6 overexpression (Dox versus control), ranked by the DESeq2 Wald statistic ^72^, and tested against the top 50 marker genes for each annotated PCW3 human embryo scRNA-seq population from Zeng et al. ^24^. Ridgeplots show the distribution of DESeq2 Wald statistics for genes within each population marker set along the ranked gene list; populations are ordered by normalised enrichment score (NES), and colour denotes FDR-adjusted p-values. (F) UMAP of scRNA-seq data from PCW2 (CS7) human embryo from Tyser et al. ^23^, coloured by cell type. (G) UMAP of TBX6 expression in the Tyser et al. dataset. (H) Projection of TBX6-upregulated gene module onto the Tyser et al. UMAP. The module was defined as the top 500 genes upregulated upon TBX6 overexpression in hTLS (adjusted p-value < 0.05, log2 fold change > 2) and filtered for overlap between the Tyser and Zeng datasets (221 genes total remaining). (I) Violin plot showing enrichment of the TBX6-upregulated gene module across cell types in the Tyser et al. dataset, using cluster annotations as in (E). (J) UMAP of scRNA-seq data from PCW3 (CS10) human embryo dataset from Zeng et al. ^24^, coloured by cell type. (K) UMAP of TBX6 expression in the Zeng et al. dataset. (L) Projection of the TBX6-upregulated gene module (as in H) onto the Zeng et al. dataset. (M) Violin plot showing enrichment of the TBX6-upregulated gene module across cell types in the Zeng et al. dataset, using cluster annotations as in (J).

**Supplementary Figure 4.**
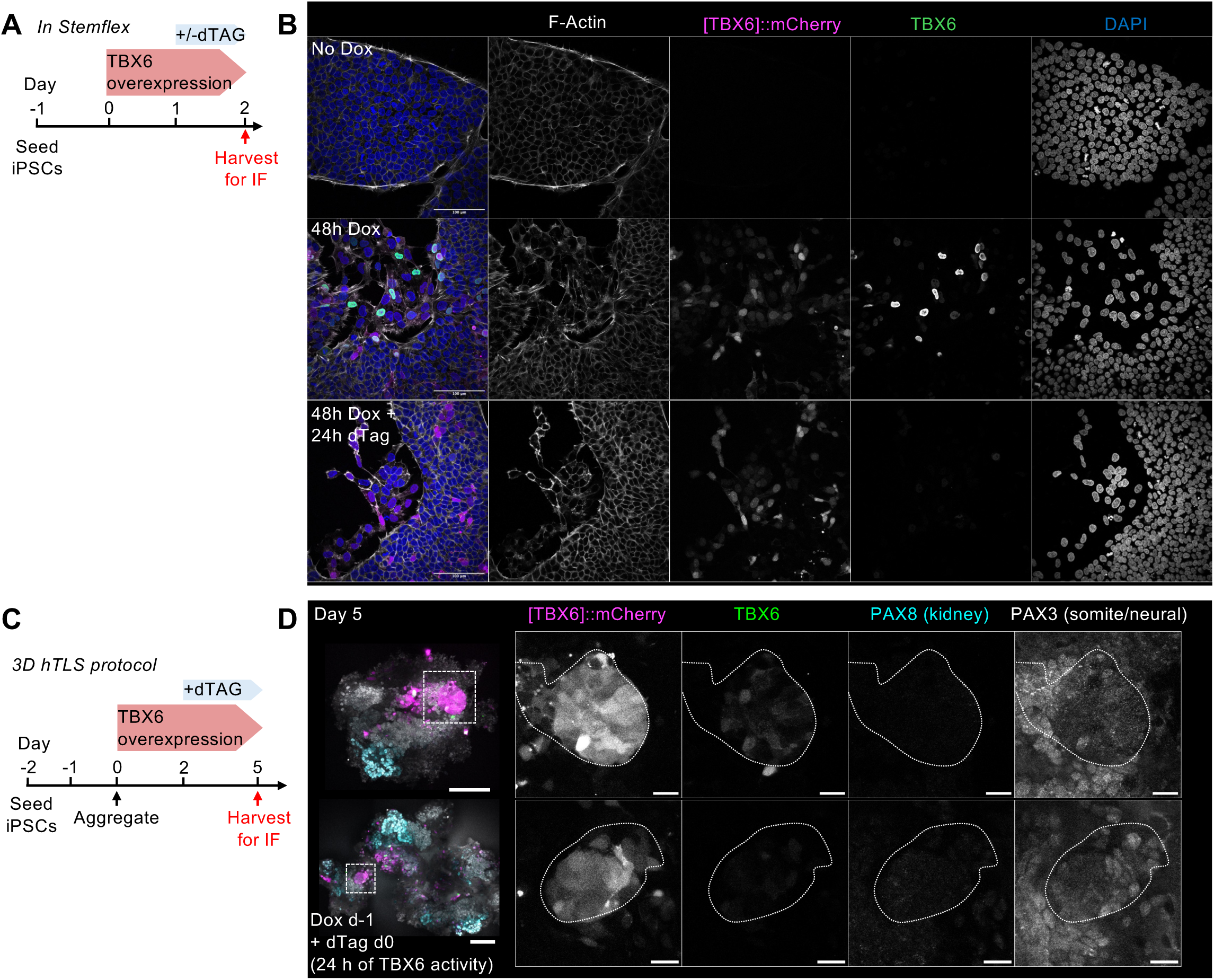
DegTrace (TBX6-FKBP-V-T2A-mCherry lineage tracing) system (related to Figure 4). (A) Experimental design to test depletion of the TBX6-FKBP fusion protein following dTAG treatment in PB-Dox-TBX6 iPSCs cultured in iPSC maintenance medium (Stemflex). (B) Immunofluorescence of F-Actin (phalloidin, grey), mCherry (magenta) and TBX6 (green), with nuclei counterstained with DAPI (blue), in representative PB-Dox-TBX6 iPSCs cultured as in (A) and either untreated (no Dox, top row), treated with Dox for 48 h (0.1 µM, middle row), or treated with Dox for 48 h followed by dTAG for 24 h (2 µM, bottom row). Scale bars, 100 µm. (C) Experimental design to test depletion of the TBX6-FKBP fusion protein in hTLS generated from PB-Dox-TBX6 iPSCs, followed by immunofluorescence (IF) analysis. (D) Immunofluorescence of mCherry (magenta), TBX6 (green), PAX8 (cyan), and PAX3 (grey) in representative day 5 PB-Dox-TBX6 hTLS treated as in (C), with Dox (0.1 µM) from day -1 followed by dTAG (2 µM) from day 0. White boxes indicate mCherry^+^ epithelial structures shown at higher magnification; dotted lines in the insets outline the mCherry^+^ structures. The top row shows an hTLS retaining low levels of TBX6 within the mCherry^+^ structure, whereas the bottom row shows an mCherry^+^ structure with TBX6 fully depleted. Scale bars: main, 100 µm; insets, 20 µm.

**Supplementary Figure 5.**
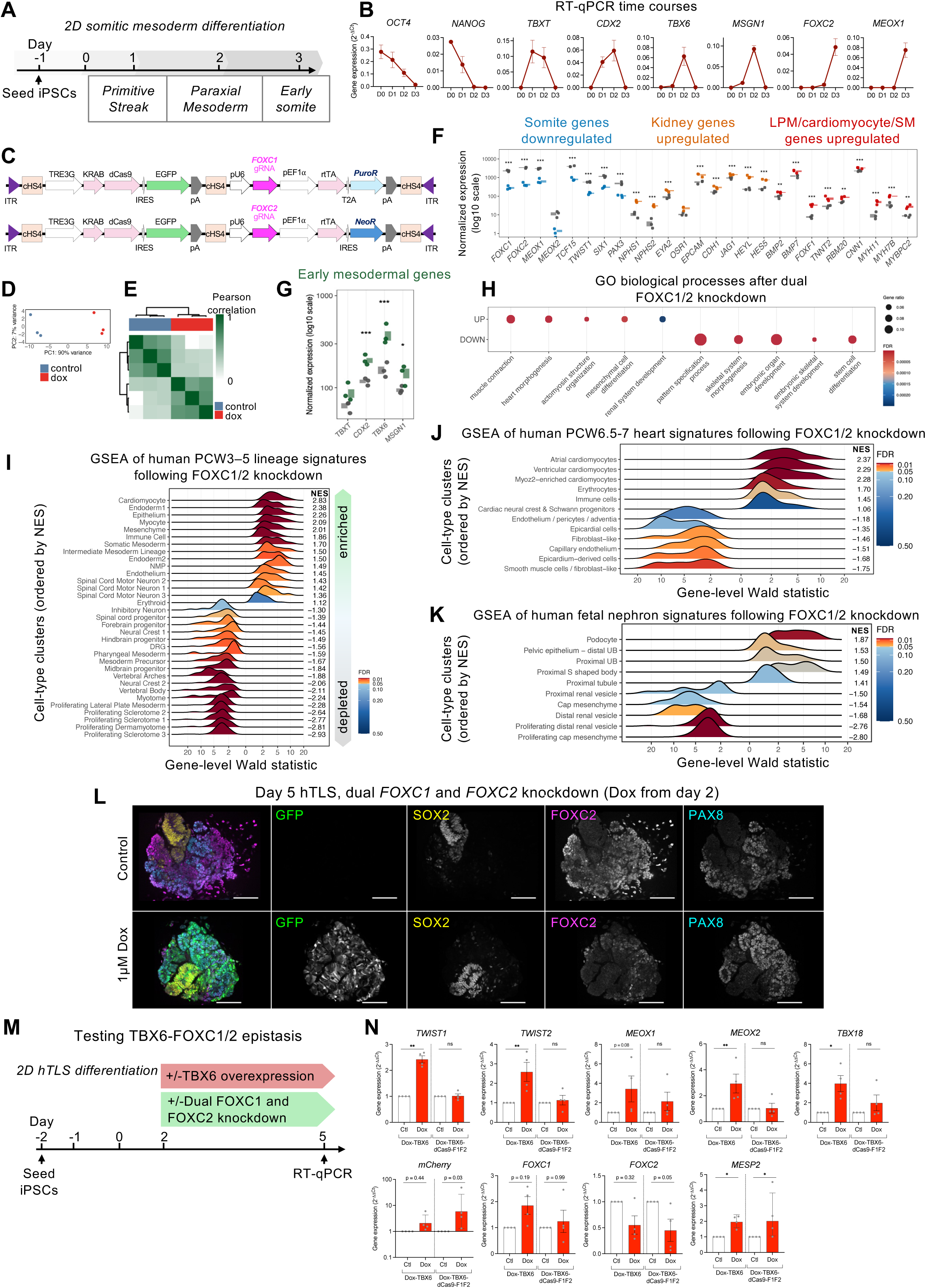
Dual FOXC1 and FOXC2 knockdown quality control and additional analyses (related to Figure 5) (A) Schematic of the 2D somitic mesoderm differentiation protocol, adapted from Loh et al. ^45^. (B) Time-course RT-qPCR analysis of selected marker genes during differentiation. Dots represent n = 2 independent differentiation batches per group, bars represent mean ± SEM. (C) Schematic of piggyBac-based constructs for Dox-inducible dual CRISPRi-mediated knockdown of *FOXC1* and *FOXC2*. (D) Principal component analysis of the dual FOXC1 and FOXC2 knockdown bulk RNA-seq dataset. (E) Sample-to-sample correlation heatmap of the dual FOXC1 and FOXC2 knockdown bulk RNA-seq dataset. (F, G) Gene-level boxplots showing DESeq2 ^72^ normalised counts (log10 scale) for (F) representative genes differentially expressed during somite differentiation including downregulated canonical somite genes (blue), upregulated kidney-associated genes (orange), and upregulated genes associated with lateral plate (LP), cardiomyocyte, or smooth muscle (SM) lineages (red), and (G) early and paraxial mesodermal markers, following dual FOXC1 and FOXC2 knockdown. FOXC1 and FOXC2 were among the most strongly downregulated genes (log2FC ≈ −3; FDR < 10⁻¹OO), confirming efficient repression. Dots represent biological replicates; boxplots indicate the IQR across replicates; grey indicates control and coloured indicates Dox treatment. Asterisks denote DESeq2 adjusted p-values (* p < 0.05, ** p < 0.01, *** p < 0.001). (H) Dot plot showing selected enriched gene ontology (GO) biological process (BP) terms among genes upregulated (UP) or downregulated (DOWN) following dual FOXC1/FOXC2 knockdown. Dot size indicates gene ratio; colour denotes FDR-adjusted p-value. (I-K) Ridge plots of gene set enrichment analysis (GSEA) of genes following dual FOXC1 and FOXC2 knockdown (Dox versus control), ranked by the DESeq2 Wald statistic ^72^, tested against marker gene sets for (I) PCW3-5 human embryo scRNA-seq populations from Zeng et al. ^24^, (J) fetal human nephron cell types from Stewart et al. ^47^, and (K) PCW6.5-7 human heart cell types from Asp et al. ^46^. Ridgeplots show the distribution of DESeq2 Wald statistics for genes within each population marker set along the ranked gene list; populations are ordered by normalised enrichment score (NES), and colour denotes FDR-adjusted p-values. (L) Immunofluorescence of hTLS generated from FOXC1/FOXC2-dCas9-KRAB iPSCs and harvested at day 5, either untreated (control, top) or treated with Dox from day -1 (1 µM, bottom), showing GFP (green), SOX2 (yellow), FOXC2 (magenta) and PAX8 (cyan), illustrating depletion of FOXC2 upon dual FOXC1 and FOXC2 knockdown. Scale bars, 100 µm. (M) Schematic of experimental design to test epistatic relationship between TBX6 and FOXC1/2 in 2D-cultured hTLS. (N) RT-qPCR analysis of somite markers in 2D-cultured hTLS derived from PB-Dox-TBX6 or Dox-TBX6-dCas9-FOXC1/2 iPSCs at day 5, either untreated (control) or treated with Dox (0.1 µM) from day -1. Dots represent mean of technical replicates from n = 4 independent differentiation batches per condition. Two-way ANOVA on log-transformed values (ΔCt), with treatment as the column factor and differentiation batch as the row factor, followed by post hoc comparisons of Dox-treated versus control conditions within each cell line. P values correspond to main treatment effect (* p < 0.05, ** p < 0.01). Data are plotted as 2^-ΔΔCt^ for visualisation, normalised to the untreated control condition within each iPSC line; bars represent mean ± SEM. ITR, inverted terminal repeat; TRE3G, third-generation tetracycline-responsive promoter; cHS4, chicken hypersensitive site 4 insulator; pA, polyadenylation signal; PuroR, puromycin resistance gene.

**Supplementary Figure 6.**
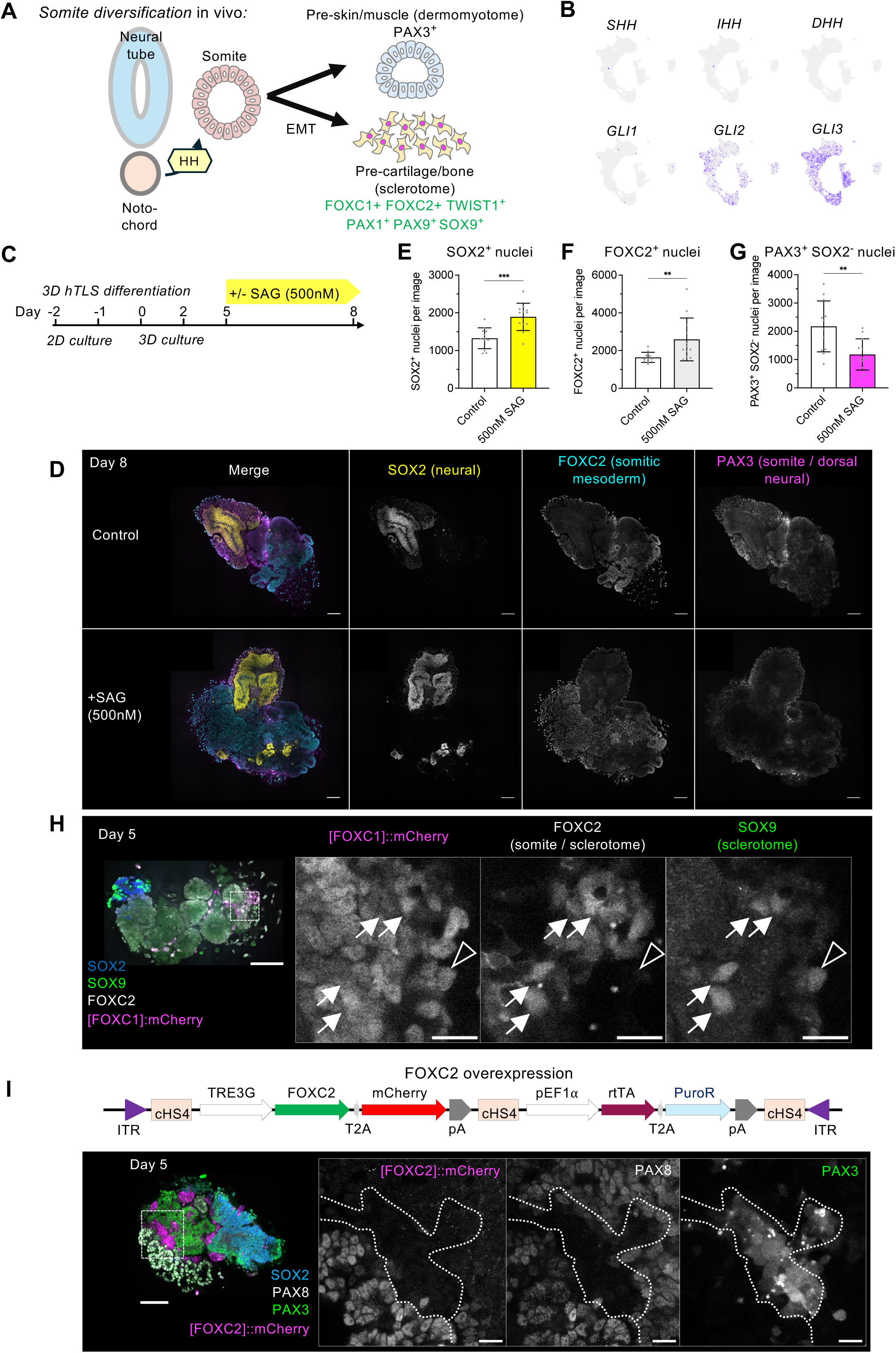
hTLS retain sclerotome differentiation potential (related to Figure 6) (A) Schematic of sclerotome differentiation *in vivo*. Hedgehog ligands secreted by the notochord signal to the ventral domain of epithelial somites to activate sclerotome-associated genes including FOXC1, FOXC2, TWIST1, PAX1, PAX9, and SOX9, to downregulate PAX3, and to initiate epithelial-to-mesenchymal transition (EMT), while dorsal somites remain PAX3^+^. (B) UMAP plots of 13,618 hTLS snRNA-seq cells (as in Figure 1E) showing minimal expression of Hedgehog ligands (*SHH*, *IHH*, *DHH*), and expression of downstream Hedgehog pathway mediators (*GLI1*, *GLI2, GLI3*). (C) Experimental design to test the effect of Hedgehog pathway activation using smoothened agonist (SAG; 500 nM) in hTLS from day 5 to day 8 of the protocol. (D) Immunofluorescence of SOX2 (yellow), FOXC1 (grey), and PAX9 (cyan) expression in day 8 hTLS cultured as in (C), either untreated (control, top row) or treated with SAG from day 5 (500 nM, bottom row). Scale bars, 100 µm. (E-G) Quantification of the number of nuclei per confocal image positive for (E) SOX2, (F) FOXC2, or (G) PAX3 (whilst also SOX2^-^) in day 8 hTLS either untreated (control, white bars) or treated with SAG from day 5 (500 nM, coloured bars). Dots represent individual hTLS (n = 12 hTLS across 2 independent differentiation batches); bars represent median ± IQR; Mann-Whitney test (** p < 0.01, *** p < 0.001). (H) Immunofluorescence of mCherry (magenta), FOXC2 (grey), and SOX9 (green) expression in day 5 hTLS generated from PB-Dox-FOXC1 iPSCs treated with Dox from day 2 (0.1 µM). White box indicates magnified region; white arrows indicate mCherry^+^ FOXC2^+^ cells expressing SOX9; arrowheads indicate a rare mCherry^-^ FOXC2^+^ SOX9^+^ cell. Scale bars: main, 100 µm; insets, 20 µm. (I) Schematic of the Dox-inducible FOXC2-overexpression strategy and corresponding immunofluorescence of SOX2 (blue), mCherry (magenta), PAX8 (grey), and PAX3 (green) expression in a day 5 hTLS generated from PB-Dox-FOXC2 iPSCs and treated with Dox from day 2 (0.1 µM). White box indicates magnified region; the dotted lines outline an mCherry^+^, PAX8^-^, PAX3^-^ cell cluster positioned between PAX8^+^ and PAX3^+^ tissue. Scale bars: main, 100 µm; insets, 20 µm.

**Supplementary Figure 7.**
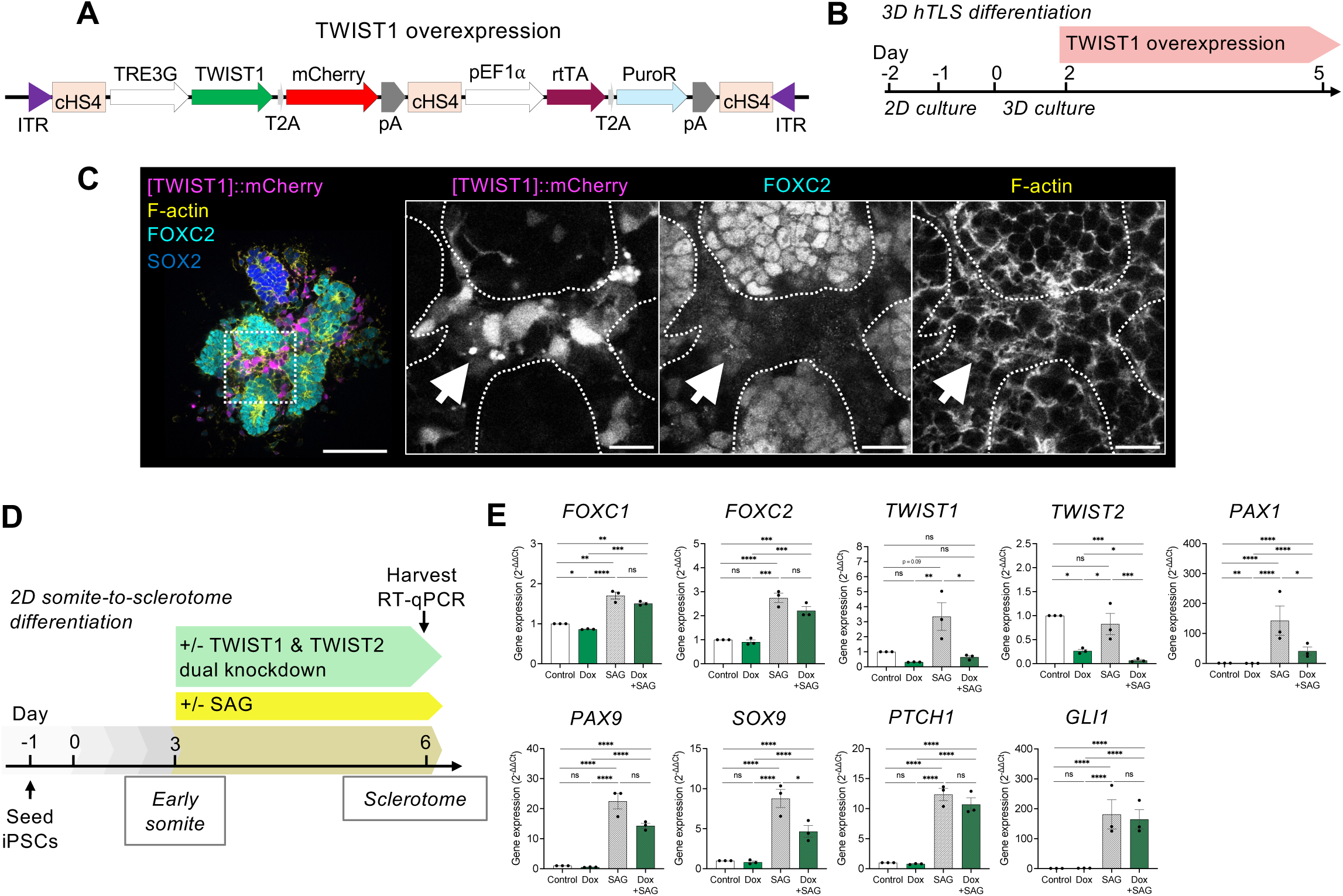
Dual TWIST1 and TWIST2 knockdown in sclerotome differentiation (related to Figure 7) (A) Schematic of a piggyBac-insertable construct for Dox-inducible overexpression of TWIST1. (B) Experimental design to test TWIST1 overexpression in hTLS. (C) Immunofluorescence of hTLS generated from PB-Dox-TWIST1 iPSCs showing mCherry (magenta), FOXC2 (cyan), F-actin (phalloidin, yellow) and SOX2 (blue) expression at day 5 following Dox treatment from day 2. The white box indicates magnified region. Dotted white lines in magnified panels indicate FOXC2^+^ epithelial structures, and white arrows indicate mCherry^+^ TWIST1-overexpressing cells occupying a subepithelial position. Scale bars: main, 100 µm; insets, 20 µm. (D) Experimental design to test dual TWIST1 and TWIST2 knockdown in a 2D sclerotome differentiation protocol (adapted from Loh et al., ^45^), followed by RT-qPCR. (E) RT-qPCR analysis of *FOXC1*, *FOXC2*, *TWIST1*, *TWIST2*, *PAX9*, *PAX1*, *SOX9, PTCH1* and *GLI1* expression in TWIST1/2-dCas9-KRAB iPSCs differentiated to sclerotome and either untreated (control), or treated from day 3 with Dox (1 µM), SAG, or combined Dox and SAG treatment. Dots represent mean of technical replicates from n = 3 independent differentiation batches per condition. Two-way ANOVA on log-transformed values (ΔCt), with treatment as the column factor and differentiation batch as the row factor. P values correspond to main treatment effect (* p < 0.05, ** p < 0.01, *** p < 0.001, **** p < 0.0001). Data are plotted as 2^-ΔΔCt^ for visualisation, normalised to the untreated control condition within each iPSC line; bars represent mean ± SEM.

## STAR METHODS

### EXPERIMENTAL MODEL AND STUDY PARTICIPANT DETAILS

#### Cell Lines

Human iPSCs were cultured as previously described with modifications ^74^. Human iPSCs used in this study were kindly gifted by Austin Smith ^75^. iPS(IMR90)-4 cells were obtained from WiCell. All iPSCs were maintained in StemFlex (ThermoFisher, A3349401) containing Pen/Strep on Geltrex (Gibco, A14133-01)-coated plates. Cells were passaged every 3-4 days using PBS with 0.5µM EDTA and re-seeded in StemFlex containing Pen/Strep and 10 µM ROCK inhibitor Y27623 (ROCKi; Tocris, 1254). Transgenic iPSC lines (TBX6-dCas9-KRAB, FOXC1/2-dCas9-KRAB, TWIST1/2-dCas9-KRAB, PB-Dox-FOXC1, PB-Dox-FOXC1[ΔFKH], PB-Dox-TBX6, PB-Dox-FOXC2, PB-Dox-TWIST1 and PB-Dox-mCherry), generated as described below, were maintained in StemFlex containing Pen/Strep, 0.5 µg/mL puromycin and/or 200 µg/mL G418 as appropriate. All cell cultures were incubated at 37°C with 7% CO_2_ and 5% O_2_.

#### Ethical Statement about use of human embryoid models

The human trunk-like structure system used in this study is a human pluripotent stem cell–derived model that although reproduces features of early post-gastrulation human developmental stages, does not generate extraembryonic tissues, and is therefore classified as a *non-integrated embryo model* under the terminology established in the 2021 International Society for Stem Cell Research (ISSCR) guidelines ^76^. In addition, the model lacks multiple embryonic lineages including, but not limited to, all anterior neural tissues, lateral mesoderm, notochord, endoderm, and non-neural ectoderm. No human embryos were used in this research. This research was conducted in accordance with the 2021 ISSCR Guidelines.

## METHOD DETAILS

### Plasmid constructs

Target gene knockdown in iPSCs was performed as previously described with modifications ^74^. pPB-Ins-U6p-sgRNAentry-EF1Ap-TetOn3G-IRES-Neo (Addgene plasmid # 183411; RRID:Addgene 183411) and pPB-Ins-TRE3Gp-KRAB-dCas9-ecDHFR-IRES-GFP-EF1Ap-Puro (Addgene plasmid # 183410; RRID:Addgene 183410) were kind gifts from Azim Surani. The SOX2:mNeon iPSC line was generated with the following plasmids: pCAS9-mCherry-Frame +0 ^77^ (gift from Veit Hornung, Addgene plasmid # 66939 ; http://n2t.net/addgene:66939 ; RRID:Addgene_66939), pCRISPaint-mNeon-PuroR [M1G] ^78^ (gift from Michael Hoelzel, Addgene plasmid # 171800 ; http://n2t.net/addgene:171800 ; RRID:Addgene_171800), and pSPgRNA ^79^ (gift from Charles Gersbach, Addgene plasmid # 47108 ; http://n2t.net/addgene:47108 ; RRID:Addgene_47108) containing gRNA targeting the human SOX2 locus (TGCCCCTCTCACACATGTGA). All gRNA oligos used were ordered using the custom DNA oligo synthesis service from Merck. Specific gRNA sequences for dCas9-KRAB mediated gene repression were designed by identifying putative transcription start sites (TSSs) for target genes according to FANTOM CAGE data publicly available on ZENBU ^80^, then the Wellcome Sanger Institute Genome Editing (WGE) CRISPR Finder ^81^ was used to identify CRISPR sequences 50-100 bp downstream of a putative TSS for optimal gene knockdown ^82^. Two gRNA sequences were tested per TSS. For generation of gRNA expression vectors against TBX6, complementary gRNA oligos were annealed and ligated into pPB-Ins-U6p-sgRNAentry-EF1Ap-TetOn3G-IRES-Neo (Addgene plasmid, 183411) by Esp3I digestion (NEB, R0734) followed by ligation with T4 DNA ligase (New England Biolabs, M0202S). Piggybac-based Dox-inducible transgene overexpression plasmids were created by first PCR amplifying (with Cloneamp Takara, 639298) the Dox-response element from pPB-Ins-TRE3Gp-KRAB-dCas9-ecDHFR-IRES-GFP-EF1Ap-Puro (Addgene plasmid # 183410) and the gRNA-rtTA expression cassette from pPB-Ins-U6p-sgRNAentry-EF1Ap-TetOn3G-IRES-Neo (Addgene plasmid, 183411), and cloning into VB240515-1632gka backbone (Vectorbuilder) by digesting with EcoRV (NEB, R3195) and PacI (NEB, R0547) and inserting using In Fusion Snap Assembly master mix (Takara, 638948). This yielded Dox-inducible dCas9-KRAB vectors that were used for knockdown of FOXC1/2 and TWIST1/2 factors; versions with Puro and Neo resistance were generated to combine 2 distinct gRNA-expressing plasmids into the same cells. For Dox-inducible expression of transgenes of interest, the dCas9-KRAB-U6-gRNA cassette was excised with NotI (NEB, R3189) and AsiSI (NEB, R0630) and transgenes were cloned in by In Fusion. FOXC1 and TWIST1 transgenes were synthesised by Twist Biosciences. mCherry was synthesised by Vectorbuilder (VB240515-1632gka). TBX6 was PCR amplified from TFORF1979 (Addgene 142303, kind gift from Feng Zhang). FKBP-V was PCR amplified from pBS_GGSG_Fkbp_2xHA_BB2 (Addgene 235503, kind gift from Brian Hendrich). PCR amplified fragments were gel extracted using Monarch gel extraction kit (NEB, T1120). Stellar competent E. coli (Takara, 636763) were used for transformation. Successful transformants were purified using QIAprep Spin Miniprep Kit (Qiagen, 27104) or QIAprep Plasmid Plus Midiprep Kit (Qiagen, 12943), then validated with Nanopore sequencing (Oxford Nanopore Technologies) by Plasmidsaurus.

### Quantitative reverse transcriptase PCR (RT-qPCR)

To isolate RNA for RT-qPCR analysis, cells were washed twice with PBS and then lysed in 350 µl RLT lysis buffer prior to RNA extraction using the RNeasy mini kit (Qiagen, 74106) with on-column DNase digestion (Qiagen, 79254) and reverse transcribed to cDNA using the QuantiTect Reverse Transcription kit (Qiagen, 205314). Quantitative PCRs were performed using PowerUp SYBR Green Master Mix (Applied Biosystems, A25742) with forward and reverse primers diluted to final concentration of 1 µM per reaction and run either on the Quantstudio 5 384-well qPCR machine (Applied Biosystems) or the StepOne Plus with 96-well block (Applied Biosystems). Statistical analyses were performed on ΔCt values (Ct_target − Ct_reference). Data were converted to 2^−ΔΔCt^ values for visualisation ^83^. Details of statistical tests used are described in the figure legends. RT-qPCR data were analysed using Prism (Graphpad, v11.0.0).

### Transgenic cell lines

TBX6-dCas9-KRAB, FOXC1/2-dCas9-KRAB, TWIST1/2-dCas9-KRAB, PB-Dox-FOXC1, PB-Dox-FOXC1[ΔFKH], PB-Dox-TBX6, PB-Dox-FOXC2, PB-Dox-TWIST1 and PB-Dox-mCherry transgenic lines were established as previously described with modifications ^74^. 6.5x10^5^ iPSCs were seeded into Geltrex-coated 6-well plates in StemFlex with ROCKi, and the following day were washed with PBS and incubated in 1ml Optimem media (Gibco, 31985070) with Revitacell supplement (Fisher, 15317447). For each well of a 6-well plate, cells were transfected with Super piggyBac Transposase expression vector (Stratech, PB210PA-1-SBI) (500 ng), together with the donor plasmid(s) to a total of 2 µg plasmid DNA mixed in 3.5 µl Lipofectamine Stem transfection reagent (Life Technologies, STEM00003) in 400 µl Opti-MEM with Revitacell. DNA-lipofectamine mixtures were incubated at room temperature for 20 minutes and then added dropwise to cells, which were then incubated at 37°C for 4 hours. Media were then replaced with 3 ml StemFlex with Revitacell. The following day, media were refreshed with StemFlex with Revitacell. 48-72 hours later, media were refreshed with fresh StemFlex with Revitacell containing antibiotics 0.5 µg/mL puromycin and/or 200 µg/mL G418 for selection. Media were refreshed with selection media every 2-3 days thereafter. A week later, surviving colonies were pooled, and knockdown efficiency was assessed by addition of 1µM Dox (a concentration we found in previous studies to be optimal ^74^) followed by RT-qPCR with primers specific to the target gene.

To make the SOX2:mNeon CRISPR-edited iPSC line, iPSCs were transfected as described above, using the following plasmids: pCAS9-mCherry-Frame +0 ^77^, pCRISPaint-mNeon-PuroR [M1G] ^78^, and pSPgRNA ^79^ containing gRNA targeting the human SOX2 locus (TGCCCCTCTCACACATGTGA). 72 hours after transfection, cells were selected with 0.5 µg/mL puromycin, and successful colonies were isolated, with mNeon expression confirmed by fluorescence microscopy.

### Human trunk-like structure generation

Human trunk-like structures (hTLS) were generated using a previously described protocol with modifications ^27^. Human iPSCs were grown to 70% confluency, washed with PBS containing 0.5µM EDTA and dissociated with Accutase (Day -2), counted using Countess II (Life Technologies, AMQAX1000), then seeded evenly onto Geltrex-coated 6-well plates at 400,000 cells per well in 2ml Nutristem hPSC XF medium (Sartorius, 05-100-1A) with ROCKi. The following day (Day -1), cells were incubated in Nutristem with 6µM CHIR (ABCR, AB253776). The next day (Day 0), cells were washed with PBS with EDTA, then dissociated with Accutase, centrifuged at 300g and resuspended in Essential 6 media (E6; Thermo, A1516401) containing 3µM CHIR and 500nM retinoic acid (RA), and counted. For 3D hTLS, 400 cells were seeded in 50µl of E6 with 3µM CHIR and 500nM RA per well of a round-bottom low-adhesion 96 well plate (Corning, CLS7007); the plate was then centrifuged at 200g for 2 minutes and incubated at 37°C to promote aggregation. By the following day (day 1) cell aggregates were visible by light microscopy at 4x magnification, at which point 150 µl E6 was added to each well gently to avoid disrupting the aggregates. On day 2, 150 µl media was carefully removed from each well, with care taken to avoid aspirating the aggregates, and 150 µl E6 containing 10% Matrigel (Corning, 356231) and 10 nM RA was gently added to each well, with Dox added as indicated. Plates were incubated at 37°C until Day 5, by which point the aggregates had elongated into trunk-like structures. To harvest for immunofluorescence analysis, structures were gently aspirated using a glass Pasteur pipette and fixed in 3% PFA for 30 minutes as described below. Alternatively, if the experiment was continued, 50 µl E6 was added to each well together with any indicated treatment, and hTLS were cultured further until harvest. For 2D hTLS, on day 0 after dissociation and counting, 4.8x10^4^ cells were seeded into Geltrex-coated 24-well plates in 500 µl of E6 with 3µM CHIR and 500nM RA per well. The following day (day 1) 375 µl medium was removed from each well and replaced with 375 µl E6 only. On day 2, 375 µl medium was removed and replaced with 375 µl E6 containing 10% Matrigel and 10 nM RA. Plates were incubated at 37°C, then on day 5 wells were washed twice with PBS then lysed for RNA extraction. The 2D hTLS protocol mirrors the 3D protocol in terms of treatment conditions with the exception that instead of growing as individual 3D structures, the 2D hTLS adhere to the 24 well plate and interconnect. Both protocols show similar marker gene expression dynamics, with the emergence of neural (SOX2^+^) cells, somitic mesoderm (FOXC2^+^) and renal mesoderm (PAX8^+^) identities.

### Immunofluorescence analysis

To harvest for immunofluorescence (IF) analysis, hTLS were transferred using a glass Pasteur pipette into a 30 ml sterilin, were centrifuged at 200g for 1 minute, washed once with PBS, then fixed in 3% PFA for 30 minutes at room temperature with gentle rotation, and then washed in PBS (3x10 min) and stored in PBS with 0.01% sodium azide at 4°C. Prior to antibody staining, samples were incubated in blocking buffer overnight at 4°C (PBS containing 0.25% Triton X-100, 5% BSA and 0.01% sodium azide). Samples were incubated in primary antibodies diluted in wash buffer (PBS containing 0.1% Triton X-100, 0.1% BSA and 0.01% sodium azide) at the indicated concentrations overnight at 4°C. Next, samples were washed in wash buffer (3x15 min at room temperature) and incubated in secondary antibodies, with DAPI (abcam, ab228549) where indicated, diluted in wash buffer at the indicated concentrations overnight at 4°C. The following day, samples were washed once in wash buffer and mounted on glass slides in Vectorshield (Vector Labs, H1000) in 120 µm spacers (Invitrogen, S24735) with coverslips overlaid.

### Imaging and image analysis

Brightfield images were taken using an EVOS M5000 microscope (Life Technologies, AMF5000SV). Confocal images were taken using a Zeiss LSM 710 or 880. Image files were analysed using Zeiss Zen (3.12) or Fiji (Image J). All immunofluorescence images presented are representative confocal slices from at least 2 independent hTLS differentiation batches. To quantify number of cells expressing a given marker, single confocal images were analysed in CellProfiler using a custom analysis pipeline ^84^. Briefly, positively stained nuclei were identified using a thresholding and declumping module (IdentifyPrimaryObject module with Otsu two-class adaptive thresholding). Smoothing factors were determined empirically. Objects (nuclei) were filtered by size, and the number of objects within that channel was measured.

### 2D somitic mesoderm differentiation

Human iPSCs were differentiated to somitic mesoderm using a previously described protocol with modifications ^45^. Human iPSCs were grown to 70% confluency, washed with PBS containing 0.5µM EDTA and dissociated with Accutase (day -1), counted using Countess II (Life Technologies, AMQAX1000), then seeded evenly onto Geltrex-coated 24-well plates at 50,000 cells per well in 500 µl Nutristem hPSC XF medium (Sartorius, 05-100-1A) with ROCKi. The following day (day 0), cells were incubated in Nutristem with 6µM CHIR (ABCR, AB253776) and 100 nM PIK90 (Tocris, 7902). Recombinant FGF2 and Activin A were omitted as Nutristem contains FGF2 and TGFß activity. The next day (day 1, primitive streak stage) media were refreshed with E6 containing 3 µM CHIR, 20 ng/ml FGF2, 1 µM A-83-01 (Tocris 2939) and 250 nM LDN 193189 (Tocris 6053). On day 2 (paraxial mesoderm stage), media were refreshed with E6 containing 1 µM A-83-01, 250 nM LDN 193189, 1 µM XAV 939 (Tocris 3748) and 500 nM PD0325.

On day 3 cells were early somite stage, at which point cells were harvested for RNA extraction of bulk RNA-seq. Alternatively, for sclerotome differentiation cells media were refreshed with E6 containing 500 nM SAG (Tocris 4366) and cells were cultured for a further 72 hours. For dCas9-KRAB knockdown experiments, Dox (1 µM) was added either at day 1 or day 3 as indicated.

### Antibodies

Primary antibodies used for protein detection, with their corresponding dilutions for immunofluorescence (IF) were as follows: rabbit anti-PAX8 [EPR13511] (Abcam, ab189249, 1:200), sheep anti-FOXC2 (R&D #AF6989, 1:200), rat anti-SOX2 (Invitrogen #14-9811-82, 1:200), goat anti-DACH2 (R&D #AF5230, 1:100), rabbit anti-ZO1 (Cell Signaling #13663, 1:200), rabbit anti-PAX6 (abcam #195045, 1:200), goat anti-TCF-2/ HNF-1 beta (R&D Systems, AF3330, 1:200), mouse anti-E-Cadherin (BD Bioscience, 610181, 1:200), mouse anti-PAX3 (R&D #MAB2457, 1:100), mouse anti-PAX3 (DSHB #PAX3-c, 1:100), rabbit anti-SOX9 (Sigma #AB5535, 1:200), goat anti-PECAM-1 (R&D #AF3628, 1:200), goat anti-TBX6 (R&D #AF4744, 1:200), rabbit anti-TBXT (Cell Signaling #81694, 1:200), Phalloidin-647 (Thermo #A22287, 1:400), RFP-568 booster (Proteintech #rb2AF568-50, 1:100), and mouse anti-Twist1 (abcam #ab50887, 1:100). Alexa Fluor 405-, 488-, 568- and 647-conjugated secondary antibodies (Invitrogen) were used for detection of primary antibodies.

### Bulk RNA-sequencing (RNA-seq)

Bulk RNA-seq of TBX6 overexpression (Figures 3 and S3) was performed on n = 4 independent differentiation batches per condition. Bulk RNA-seq of dual FOXC1/2 knockdown (Figures 5 and S5) was performed on n = 3 independent differentiation batches per condition. For each sample, at least 100,000 cells per condition were collected at the indicated time points. Cells were washed with PBS containing EDTA, dissociated using Accutase, pelleted by centrifugation, and lysed in DNA/RNA Shield (Zymo, R1100-50).

Bulk RNA-seq libraries were prepared and sequenced by Plasmidsaurus using an Illumina-based 3′ end counting approach. Total RNA was extracted using a bead-based method, and mRNA was reverse transcribed using poly(dT) priming, followed by second-strand synthesis, tagmentation, and PCR amplification. Libraries incorporated unique molecular identifiers (UMIs) to enable deduplication of PCR duplicates and were sequenced as single-end reads (∼90 bp) enriched at the 3′ end of transcripts.

Sequencing reads were processed using a standardized pipeline including quality filtering, alignment to the reference genome using STAR, UMI-based deduplication, and gene-level quantification using featureCounts to generate count matrices for downstream analysis.

### Bulk RNA-seq analysis

Count matrices were imported into RStudio (version 2024.09.1+394) running R version 4.4.2, and differential gene expression analysis between control and Dox-treated conditions was analysed using DESeq2 on raw count data ^72^. The significance threshold for differential expression was set as log2 fold change > 0.58 (corresponding to fold change > 1.5) and adjusted p-value < 0.05. Transformed, variance-stabilised expression values were used for principal component analysis and sample-to-sample correlation heatmaps. Expression plots for individual genes used DESeq2 normalised counts. Gene ontology (GO) analysis was performed with the enrichGO function from clusterProfiler ^85^.

### Single-nucleus RNA-seq library preparation

For single-nucleus RNA-sequencing experiments, cell pellets or hTLS containing 0.5-1 M cells harvested at the indicated time points were snap frozen. Pellets were thawed and resuspended in 1 ml of ice-cold nuclei permeabilization buffer (10 mM Tris-HCl pH7.4, 10 mM NaCl, 3 mM MgCl_2_, 0.2% IGEPAL CA300, 0.1 mg/ml BSA, 1x protease inhibitor cocktail (Roche cat.11873580001), 0.5 U/µl ea Superase RNAse Inhibitor and RNAaseOUT). Cells were then permeabilized for 5 min and centrifuged at 600g for 5 min at 4°C. Cells were lightly fixed by resuspending in 475 µl fixation buffer (1xPBS, 0.2% formaldehyde, 0.5 U/µl ea Superase RNAse Inhibitor and RNAaseOUT) and incubated on ice for 5 min. Fixation was stopped by adding 2 µl 1M Tris-HCl pH7.4 containing 0.1 mg/ml BSA. Nuclei were pelleted at 600g for 5 min at 4°C and resuspended in 1xPBS containing 1x protease inhibitor cocktail and 0.5 U/µl Superase RNAse Inhibitor and 0.5 U/µl RNAaseOUT.

For in situ reverse transcription (RT) reaction 50,000 cells for each sample were aliquoted in 0.2 ml PCR tubes containing 2.5 µM sample specific barcoded primers as previously described ^86^. To each tube RT mix (1x Maxima H buffer, 0.5 mM dNTPs, 0.1 mg/ml BSA, 400U Maxima Reverse H Minus Reverse Transcriptase (Thermo Scientific) and 0.5 U/ul ea Superase RNAse Inhibitor and RNAaseOUT) was added to a final volume of 20 µl. The reverse transcription was performed in a thermocycler with the following program: Step 1: 50°C × 10 min; Step 2 (3 cycles): 8°C × 12 s, 15°C × 45 s, 20°C × 45 s, 30°C × 30 s, 42°C × 2 min, 50°C × 5 min; Step 3: 50°C × 10 min. After the reaction, the nuclei were transferred and pooled on ice into a 1.5-ml tube prewashed with 5% BSA in 1x PBS. Then Triton X-100 was added to a final percent of 0.075 and nuclei were pelleted at 800g for 5 min at 4°C.

For ligation-based combinatorial barcoding the pooled nuclei were resuspended in 1 ml of 1x NEBuffer3.1 (NEB) and transferred to 2,910 µl of ligation buffer (1x T4 ligase buffer (NEB), 0.35 mg/ml BSA, 25 mM NaCl, 40,000 U T4 ligase (NEB), and 0.25 U/µl Superase RNAse Inhibitor and 0.25 U/µl RNAaseOUT). Using a multichannel pipette, 40 µl of the ligation mix was distributed to the 96 wells of the first R02 barcode plate ^86^. The plate was sealed and placed on a Thermomixer (Eppendorf) and mixed for 30 sec at 37°C at 1200 rpm. Then incubated for a further 30 min at 37°C at 300 rpm. After incubation the plate was removed and 10 µl of R02 blocking solution (26 µM R02 blocking oligo 5’ ATCCACGTGCTTGAGAGGCCAGAGCATTCG in 1x T4 ligase buffer) was added to each well. The plate was re-sealed and mixed again for 30 sec at 1200 rpm before incubating for an additional 30 min at 37°C. The nuclei from each well were then pooled into 1.5 ml tubes and pelleted at 800g for 5 min at 4°C. The second round of ligation was then carried out using R03 biotin tagged barcode plate ^86^ similarly to the first round, except that after the 30 min ligation reaction, blocking solution (26 µM quencher oligo 5’ GTGGCCGATGTTTCGGTGCGAACTCAGACC in 0.25 M EDTA) was added to quench the reaction. After 2 rounds of ligation-based barcoding, the pooled nuclei were resuspended in 1x PBS with 0.5 U/µl Superase RNAse Inhibitor and 0.5 U/µl RNAaseOUT, counted and aliquoted into 10 µl sublibraries containing 3,000 nuclei.

For each sublibrary nuclei were lysed and cross-links reversed by adding 10 µl of lysis buffer (10 mM Tris-pH 7.5, 1 M NaCl, 0.5 mM EDTA, 2% SDS) and incubating at 65°C for 5 hrs. Biotin barcoded cDNA molecules were purified by adding 80 µl of dynabead solution (10 ul MyOne Streptavidin C1 dynabeads (Invitrogen) in 10 mM Tris-pH 7.5, 1 M NaCl, 0.5 mM EDTA) and incubating for 1 h at 24°C at 1000 rpm. Bound cDNAs were washed twice with 200 µl BW buffer (10 mM Tris-pH 7.5, 1 M NaCl, 0.5 mM EDTA) followed by 2 additional washes with 200 µl Tris-T solution (10 mM Tris-HCl pH7.5, 0.1% Tween 20). For incorporating a 5’ adaptor onto the streptavidin bound cDNA a 50 µl template switching mix (1x Maxima H buffer, 4% Ficoll, 1 mM dNTPs, 2.5 µM TS oligo (AAGCAGTGGTATCAACGCAGAGTGAATrGrG+G where rG is RNA base), 500 U Maxima H minus RT, 15U Superase RNAse Inhibitor and 20 U RNAaseOUT) was added to each sublibrary. The bead solution was then incubated at 25°C for 30 minutes and then at 42°C for 90 minutes with mixing at 1000 rpm. After the incubation beads were washed twice with 200 µl Tris-T solution.

Washed beads were resuspended in 50 µl of PCR reaction mix (1x Q5 reaction buffer, mM dNTPs, 0.5 uM ea Rev (AAGCAGTGGTATCAACGCAGAGT) and For (CAGACGTGTGCTCTTCCGATCT) primer, 1U Q5 High-Fidelity DNA polymerase (NEB)) and amplified with the following program: Step 1: 95°C × 1 min; Step 2 (6 cycles): 98°C × 30 s, 70°C × 15 s, 72°C × 3 min; Step 3: 72°C × 1 min. PCR reactions were purified using a 0.85X ratio of SPRI beads (Beckman). To determine the optimal PCR cycle to give 1/3^rd^ of a saturated signal we repeated the PCR using 1/10^th^ of the purified PCR reaction and EvaGreen (Biotium) in a qPCR machine. Using the optimal cycle we PCR amplified the remaining cDNA and purified the reactions using 0.85X ratio of SPRI beads. To normalize the amplified cDNA fragment lengths for sequencing we trimmed the libraries using Tn5 transposase (Diagenode) loaded with (5’-[Phos] CTGTCTCTTATACACATCT, 5’-TCGTCGGCAGCGTCAGATGTGTATAAGAGACAG) to a size range of 200-1000bp. Each trimmed sublibrary was then PCR amplified using a unique forward index P7Trueseq barcode oligo (5’ CAAGCAGAAGACGGCATACGAGATXXXXXXGTGACTGGAGTTCAGACGTGTGCTCT TCCGATC) and reverse oligo (AATGATACGGCGACCACCGAGATCTACACTAGATCGCTCGTCGGCAGCGTCAG).

Purified libraries were then sequenced on a NovaSeq 6000 using 100 bp paired end reads and a read depth of 25-30,000 reads per cell.

### Single-nucleus RNA-seq data preprocessing

The Paired-Tag pipeline as previously described ^87^ (https://github.com/cxzhu/Paired-Tag) was used to process raw sequencing reads. Briefly this set of scripts takes raw reads and extracts cellular barcodes from read2 and then uses read1 sequence to map reads to human genome reference hg38. Finally demultiplexed and mapped data from different sublibraries are merged to generate the final cell barcode gene count matrix.

### Single-nucleus RNA-seq analysis

Single nucleus transcriptomes were analysed from hTLS at days 2-7 across 2 independent differentiation batches, with additional day 0 and day 7 samples from independent batches, and undifferentiated SOX2:mNeon iPSC cells, included. Filtered feature-barcode matrices of snRNA-seq data were converted into Seurat objects using the Seurat R package (version 5.3.1) ^88^, merged into a single object, and quality control filtered for cells containing > 500 and < 5000 genes and < 10% mitochondrial reads. An average of ∼1,500 genes and ∼1% mitochondrial reads were detected per cell, consistent across time points (Figures S1E,F). Gene expression counts were normalised using SCTransform ^89^, followed by PCA. Visualisation and clustering were performed using the functions RunUMAP and FindNeighbors, followed by FindClusters using clustering resolution as stated in the figure legends. Clustering identified a minority of clusters composed of cells with consistently lower gene detection relative to the dataset average. These clusters lacked detectable expression of established lineage marker genes and were excluded from downstream analyses due to insufficient information for reliable cell type annotation. Mesodermal cells were isolated by clustering the full filtered dataset at low resolution (0.03; Figure S1M), and re-clustered using the top 41 principal components at a resolution of 0.5. Marker genes for each cluster were identified with Wilcoxon rank sum test in Seurat (FindMarkers). For differential expression analyses, thresholds of log2 fold change > 1 and adjusted p-value < 0.005 (or 1e-8 for selected analyses) were applied, while full results were used for visualisation in volcano plots. Cell identities were manually annotated by expression of established canonical marker genes.

### Pseudotime analysis

To identify distinct mesodermal lineages arising during hTLS development, the mesoderm subset was clustered with resolution 0.5, and Slingshot ^36^ trajectories were inferred using cluster 0 as the root and clusters 2, 1, 6 and 4 as terminal states. tradeSeq ^37^ was used to model gene expression dynamics along each trajectory and for statistical analyses. To identify genes defining terminal cell states, the tradeSeq endpoint differential expression test (diffEndTest) ^37^ was used, which tests for differential expression between lineage endpoints. For each lineage, genes were ranked using a combined endpoint significance score defined as the sum of -log10-transformed p-values from the three pairwise endpoint contrasts involving that lineage. The top 20 ranked genes for each lineage were selected for visualisation. For heatmap visualisation (Figure 1K), expression dynamics along pseudotime were obtained from the fitted tradeSeq GAM as predicted expression values and transformed using log1p. Gene expression trajectories were standardised by computing z-scores relative to the gene’s mean fitted expression across all pseudotime lineages. Heatmap values thus represent global standardised expression, with positive values (red) indicating expression above the gene’s global mean and negative values (blue) indicating expression below the mean. To reduce the visual influence of genes with low absolute expression levels, standardised values were scaled by an expression-dependent weight derived from the fitted log1p expression. Specifically, z-scores were multiplied by an expression-dependent logistic weighting function derived from the fitted log1p expression, thereby downweighting pseudotime regions with low fitted expression and therefore less likely to reflect biologically meaningful expression. Weighted z-scores were clipped to ±2.5 to limit the visual influence of extreme values.

### Motif enrichment

Motif enrichment analysis was performed using HOMER (v4.11) ^90^ on ATAC-seq peaks from early somite populations generated using the same differentiation protocol as described by Loh et al ^45^. Public ATAC-seq peaks corresponding to early somite cells (GEO accession GSM2257295) were used as the accessible chromatin reference set. Peaks located within ±50 kb of transcription start sites of genes significantly downregulated following FOXC1/2 knockdown were identified using BEDTools. These regions were tested for transcription factor motif enrichment relative to a background set consisting of ATAC-seq peaks located within ±50 kb of transcription start sites of genes expressed but not significantly altered following FOXC1/2 knockdown. Motif enrichment was calculated using HOMER (findMotifsGenome.pl) with sequences extracted from the hg19 human genome assembly. Significantly enriched motifs were ranked by hypergeometric p-value and FDR. Enriched motifs were interpreted as candidate transcriptional regulators acting at FOXC-dependent regulatory elements during early somite specification.

### Mapping scRNA-seq to reference human embryogenesis datasets

We projected our data (query dataset) on two reference datasets: Zhao et al. for the earlier time points (days 0-4) ^25^, and Zeng et al. for the later ones (days 5-7) ^24^. For the Zhao et al. reference, we used the projection pipeline provided by the authors themselves for projecting any user’s query dataset onto their reference (R function: projecting_query_dataset.R). We modified only one parameter of the standard pipeline: para.list$cor.cutoff <-0.2 instead of 0.5. We projected our datasets one sample at the time. The output of this pipeline matches each cell of the target to its closest location in the reference’s umap, which allows to represent the reference landscape as published by the authors with the query dataset projected on top. For the Zeng et al dataset, we built our own projection pipeline, which consists of two parts: integration and projection. For the integration, we used the scVI method from the scVI-tools (10.1038/s41592-018-0229-2) to correct the raw counts of the reference and target datasets. Correcting the counts was necessary due to the different chemistries of the reference and the query datasets. Once the counts were corrected, we computed 50 pcs, that were then used for projecting one sample of the query datasets at the time, following these steps: we computed the distance matrix among the pcs of the reference and those of the query dataset; for each cell in a query sample, we gave a score 1 to its 15 nearest neighbours (NN) of the reference based on the distance matrix, and 0 otherwise; we computed the adjacency (or similarity) score of each cell of the reference to each cell of the query dataset as the average of the scores 1 (cell being a NN) or 0 (cell not being a NN). As a result, we can visualize the originally published umap of the reference with our own data projected on top in terms of similarity scores.

### QUANTIFICATION AND STATISTICAL ANALYSIS

Statistical analyses of bulk RNA-seq data were performed with DESeq2. Analyses of snRNA-seq data were performed with Seurat. Marker genes were analysed with Wilcoxon rank sum test. Thresholds for volcano plots were set as |log2FoldChange| > 0.58 (±1.5-fold) and adjusted p-value < 0.05, and for heatmaps were set as log2FoldChange >1 (>2-fold) and adjusted p-value < 1e-8 (Fig S1N) and adjusted p-value < 0.005 (Fig S1P). Statistical analyses of confocal microscopy quantifications and RT-qPCR data were performed within Prism. For image quantification, data were analysed with Mann-Whitney test and visualised in plots with dots representing individual hTLS and bars represent median ± IQR. For RT-qPCR analyses, individual data points represent means of up to 4 technical replicates from a single independent differentiation batch, which were used as ‘n’s for statistical analyses. Biological replicate data were assumed to have a normal distribution, while to account for batch effects, data were analysed using two-way ANOVA with batch as row factor and treatment as column factor. RT-qPCR data are visualised in plots with bars representing mean ± SEM. Asterisks in plots represent p values as follows: * (p < 0.05), ** (p < 0.01), *** (p < 0.001), **** (p < 0.0001). All the details of quantifications and statistical analyses are fully described in the main text, figure legends, and method details section.

